# Decoding The Nuclear Genome of The Human Pathogen *Babesia duncani* Shed Light on its Virulence, Drug Susceptibility and Evolution among Apicomplexa

**DOI:** 10.1101/2022.05.09.491209

**Authors:** Stefano Lonardi, Pallavi Singh, Qihua Liang, Pratap Vydyam, Eleonora Khabirova, Tiffany Fang, Shalev Gihaz, Jose Thekkiniath, Muhammad Munshi, Steven Abel, Gayani Batugedara, Mohit Gupta, Xueqing Maggie Lu, Todd Lenz, Sakshar Chakravarty, Emmanuel Cornillot, Yangyang Hu, Wenxiu Ma, Luis Miguel Gonzalez, Sergio Sánchez, Estrella Montero, Karel Estrada, Alejandro Sánchez-Flores, Omar S. Harb, Karine G. Le Roch, Choukri Ben Mamoun

## Abstract

*Babesia* species are tick-transmitted apicomplexan pathogens and the causative agents of babesiosis, a malaria-like disease of major medical and veterinary importance. Of the different species of *Babesia* reported so far, *Babesia duncani* causes severe to lethal infection in patients. Despite the highly virulent nature of this parasite and the risk it may pose as an emerging pathogen, little is known about its biology, metabolic requirements, and pathogenesis. *B. duncani* is unique among apicomplexan parasites that infect red blood cells in that it can be continuously cultured *in vitro* in human erythrocytes but can also infect mice leading to fulminant babesiosis infection and death. Here we have taken advantage of the recent advances in the propagation of this parasite *in vitro* and *in vivo* to conduct detailed molecular, genomic and transcriptomic analyses and to gain insights into its biology. We report the assembly, 3D structure, and annotation of the nuclear genome of this parasite as well as its transcriptomic profile and an atlas of its metabolism during its intraerythrocytic life cycle. Detailed examination of the *B. duncani* genome and comparative genomic analyses identified new classes of candidate virulence factors, suitable antigens for diagnosis of active infection, and several attractive drug targets. Translational analyses and efficacy studies identified highly potent inhibitors of *B. duncani* thus enriching the pipeline of small molecules that could be developed as effective therapies for the treatment of human babesiosis.

## INTRODUCTION

Recent reports suggest an increase in the incidence of tick-borne bacterial, parasitic and viral infections in the United States and worldwide (1). This is largely due to changes in environmental factors that led to the expansion of the geographical distribution of the tick vectors, several of which carry multiple human pathogens (1). Among these pathogens, *Babesia* parasites are considered a serious threat to humans and animals (2, 3). Clinical manifestation of babesiosis in humans includes malaria-like symptoms, disseminated intravascular coagulation, pulmonary edema and renal insufficiency (2). The mortality rate from *Babesia* infections can approach 9% in hospitalized babesiosis patients, and ∼20% in immunocompromised hosts (3, 4). The death rate from transfusion-transmitted babesiosis may be as high as 28% (5). Recommended treatment for human babesiosis consists of two drug combination, quinine and clindamycin for treatment of severe disease, and atovaquone and azithromycin for treatment of mild disease (2, 6).

All cases of human babesiosis reported in the United States have been linked to either *Babesia microti* (majority of cases), *B. duncani* or a *B. divergens*-like species called MO1 (3). Whereas *B. microti* is transmitted by *Ixodes scapularis*, which transmits the agents of Lyme disease (*Borrelia burgdorferi*) and human granulocytic anaplasmosis (*Anaplasma phagocytophilum*), available data suggest that *B. duncani* is transmitted by the hard tick *Dermacentor albipictus* (7). The first clinical isolate of *B. duncani* (WA1) was collected in 1991 from a patient who lacked all the typical risk factors for fulminating babesiosis (i.e. old age, splenectomy and weak immune system) (8). Phylogenetic analysis based on the parasite’s mitochondrial proteins revealed that *B. duncani* defines a new lineage closely related to *B. conradae*, a causative agent of canine babesiosis, and distinct from *B. microti*, *B. bovis*, *Theileria species*, *Plasmodium* species and *Toxoplasma gondii* (9). In four reported clinical cases of *duncani*-babesiosis in California, three patients recovered following treatment with a combination of clindamycin and quinine (10), whereas a fourth patient presented with high parasitemia, had a cardiopulmonary arrest and died one day after treatment, which included clindamycin, quinidine, doxycycline, pentamidine, red cell transfusion and hemodialysis (10). Recent studies using a continuous *in vitro* culture of the parasite in human red blood cells showed that *B. duncani* has low susceptibility to atovaquone (IC_50_ ∼500 nM), azithromycin (IC_50_ ∼5 µM), clindamycin (IC_50_ ∼20 µM) and quinine (IC_50_ ∼12 µM) (11). Consistent with the severe clinical outcome of *B. duncani* infection in humans (8,12–15), studies in immunocompetent hamsters, and both immunocompetent and immunocompromised mice confirmed the high virulent status of *B. duncani* as infection with this parasite results in severe disease manifestation including Respiratory Distress Syndrome accompanied with atelectasis, pneumothorax and pulmonary hemorrhage rapid increase in parasite load and spontaneous death with a very high mortality rate (>90%) (16, 17). Despite the highly virulent properties of this parasite, little is known about its biology, evolution and mechanism of virulence. Here we report the first completed nuclear genome sequence, assembly, 3D structure, and transcriptional annotation of *B. duncani*. Our analysis revealed that the parasite has evolved new classes of multigene families that based on their expression profile seem to be needed either during the intraerythrocytic cycle or following transmission into the tick. Drug target mining and *in vitro* efficacy studies identified the antifolates, pyrimethamine and WR99210, as excellent inhibitors of parasite development within human erythrocytes. The high potency of these compounds provides promise for the future development of more effective therapies.

## RESULTS

### Genome Sequencing and Assembly using PacBio HiFi reads, Bionano Genomics optical map, Illumina WGS and Illumina Hi-C

*B. duncani* asexual development in human and the reservoir mule deer, its alternative mode of transmission through blood transfusion, its sexual development in the tick vector, and its developmental stages within human red blood cells are depicted in Fig. 1. To gain further insight into the genomic content and structure of *B. duncani*, we took advantage of the recently developed continuous *in vitro* culture system of the parasite in human red blood cells (11) to purify its total genomic DNA and subjected it to whole genome sequencing. A high-quality genome assembly was produced by processing PacBio HiFi WGS long reads (∼150x coverage) and Illumina WGS paired-end reads (∼130x coverage) (Fig. S1). A Bionano optical map was also generated to correct and scaffold the assembly: the map generated using the restriction enzyme identified six molecules of sizes 4.5Mb, 2Mb, 1.4Mb, 1.1Mb, 474Kb and 451Kb, respectively, for a total of 10Mb (Table S1 **and** Fig. S2). PacBio HiFi reads were first filtered of mitochondrial and apicoplast genomes (9) to enrich for nuclear DNA sequences, then assembled with HiCANU (18). The *de novo* draft assembly produced by HiCANU consisted of 167 contigs, including four contigs over 1Mb. The total assembly length was 10.37Mb with a N50 of ∼1.3Mb, L50 of 3 contigs, and a NG50 of ∼1Mbp (**Table 1**). The draft assembly was compared with the Bionano optical map to detect possible mis-joins and to create scaffolds (Fig. S2). The Bionano Hybrid Scaffolding pipeline detected no chimeric contigs and created four scaffolds, two of which were composed by two HiCANU contigs with a 16-19Kb gap in-between them. The rest of the assembled HiCANU contigs were not scaffolded and kept for the final assembly. The scaffolded assembly was polished using Polypolish (19): two rounds of polishing using Illumina WGS reads only corrected less than 2K bases in the genome, which confirmed the very high-quality nucleotide level assemblies obtained from HiFi reads. The final polished assembly had five chromosome-level scaffolds of 3.13Mb (Chromosome I), 1.58Mb (Chromosome II), 1.42Mb (Chromosome IIII), 1.07Mb (Chromosome IV), and 0.35Mb (Chromosome V) with a total of only 35Kb in two gaps (see **Table 1** for other statistics). The assembly pipeline is illustrated in Fig. S1. Additional details of the pipeline are provided in Material and Methods.

**Figure 1:**
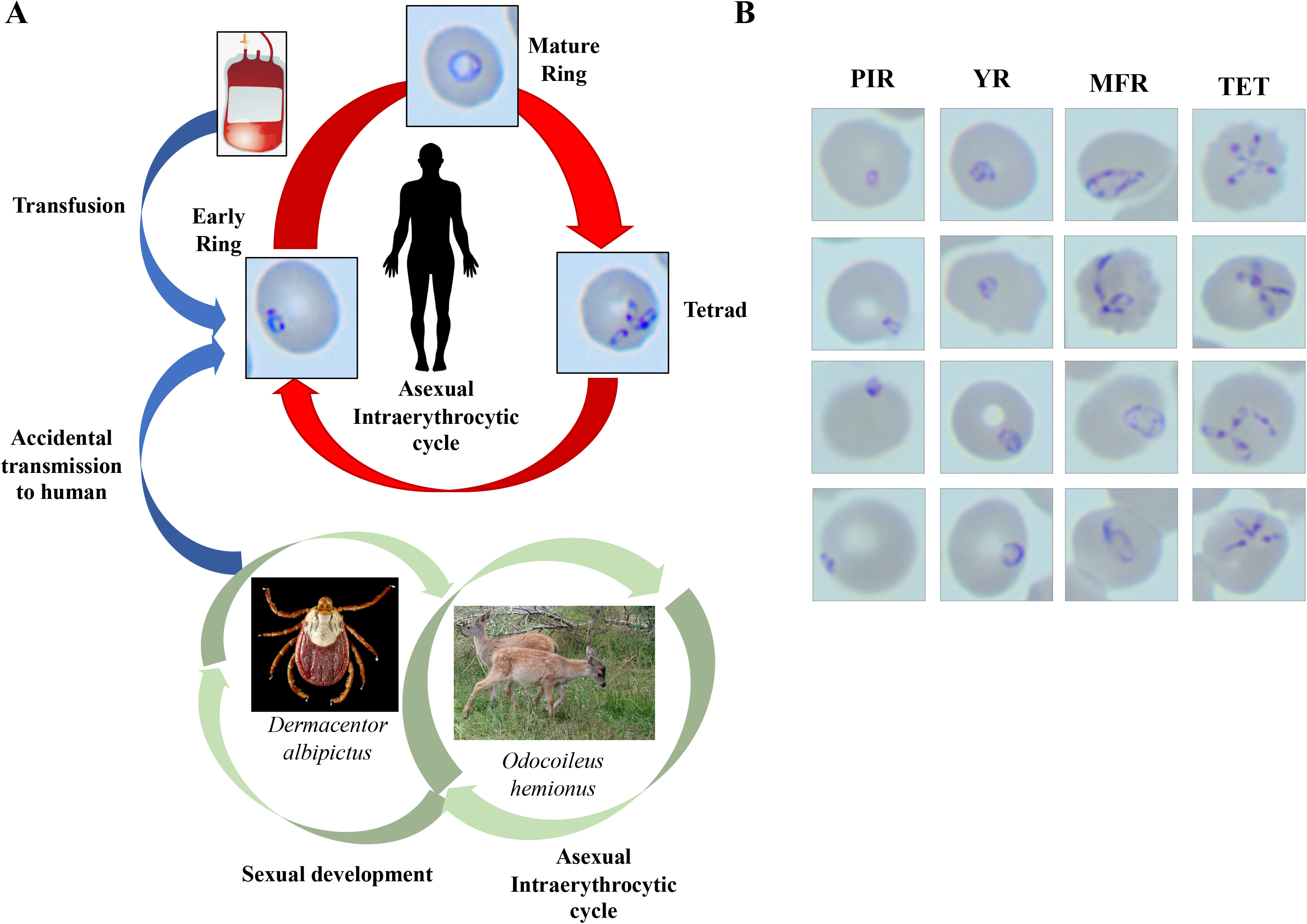
**A.** Schematic representation of the life cycle of *B. duncani* in humans, tick vector and mule deer. **B.** Representative images of the various forms of *B. duncani* grown in human erythrocytes *in vitro*. PIR: Post-invasion rings; YR: young rings; MFR: mature and filamentous rings; TET: tetrads

We also checked the assembly for mis-joints and assembly errors using the long-range chromatin contact frequency information provided by Hi-C analysis We recently used this methodology to correct the genome sequences of *Plasmodium knowlesi* (20) and *Toxoplasma gondii* (21). The contact map for *B. duncani* WA1 is shown on Fig. 2A-2B. A close examination of the contact maps indicates that the assembly has no large mis-joints or mis-assemblies, and consistent with other apicomplexan parasites (21), the centromeres interact strongly with each other. Such feature revealed that chromosome 1 has a metacentric centromere, whereas chromosomes 2 and 3 have a telocentric centromere, and chromosomes 4 and 5 have an acrocentric centromere (as illustrated in the diagram below Figure 2B). A similar organization was observed in three of the four chromosomes of *Babesia bovis* (22). Using BUSCO v5 (23), a software tool that measures completeness and redundancy of a genome in terms of expected gene content, we found that the *B. duncani* WA1 nuclear genome has 95.1% of the gene models derived from the apicomplexa_odb10 database (85.2% single-copy, and 9.9% duplicated) with another 1.8% fragmented gene models, and only 3.1% of the gene models missing. The chromosomal organization of the *B. duncani* nuclear genome was further validated using PFGE analysis. Five bands with approximate sizes of ∼3.1 Mb (Chromosome I), ∼1.81 Mb (Chromosome II), ∼1.37 Mb (Chromosome III), ∼1.05 Mb (Chromosome IV) and <1 Mb (Chromosome V) were identified using this analysis (Figure 2C). Using a specific telomeric probe from *P. berghei*, each of the 5 chromosomes was labeled using Southern blot analysis (Figure 2C). Altogether these results demonstrate that the *B. duncani* WA1 nuclear genome consists of five chromosomes. To investigate further the genome-wide chromatin organization of the *B. duncani*, a three-dimensional (3D) model was built from the Hi-C contact maps (Figure 2D). The five chromosomes of *B. duncani* showed strong interactions among the centromeres (Figure 2C and 2D), an organization similar to that of seven recently reported 3D structures of apicomplexan parasites including that of the *Babesia microti* genome (21, 24). Interaction between the telomere ends was also observed to a lesser extent leading to partial interaction between the centromeres and telomeres.

**Figure 2.**
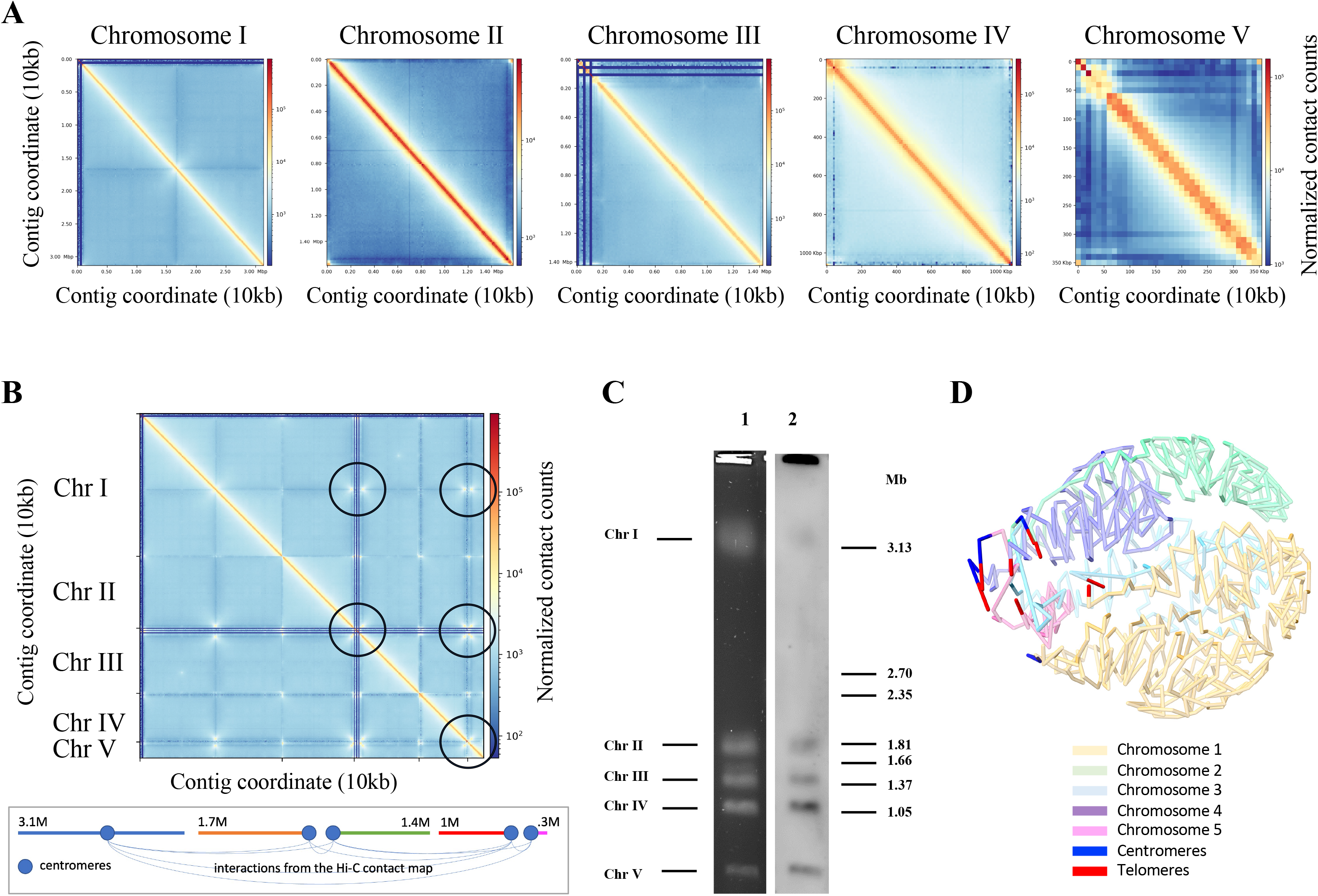
**A.** Hi-C inter-chromosomal contact maps for the five *B. duncani* chromosomes (10Kb bins) **B.** Full Hi-C contact map for *B. duncani* (10Kb bins); black circle highlight strong intra-chromosomal interaction, which are summarized in the panel below; X-shaped interactions typically indicate the presence of centromeres **C.** Pulsed field gel electrophoresis (PFGE) (line 1) and subsequent Southern-blot analyses using a *Plasmodium berghei* telomeric probe (line 2) showing the number and approximate size of *B. duncani* chromosomes (Chr) : ∼3.1 Mb, (Chr I), ∼1.81, Mb (Chr II), ∼1.37 Mb (Chr III), ∼1.05 Mb (Chr IV) and <1 Mb (Chr V). *Hansenula wingei* DNA chromosomes were used as DNA markers. The manufacture’s estimates of the sizes of chromosomes are indicated in Megabase pairs (Mb) on the left of the picture. **D.** 3D genome structure of *B. duncani* derived from the contact map interactions.

### *B. duncani* genome annotation

To accurately predict gene loci, we generated Illumina RNA-Seq and PacBio IsoSeq data using RNA isolated from parasites propagated in human red blood cells *in vitro*. Total RNA samples were processed, and cDNA generated as described in *Material and Methods*. The gene annotation pipeline used FunAnnotate and InterProScan to determine gene loci and functional annotations. The final set of gene models included 4,222 gene loci (including 52 tRNA genes) with an average length of 1,656 bp (average protein length was 499aa) (**Table I**). The pipeline identified 12,954 exons with an average length of 356 bp (**Table I**). Of the predicted 4,170 genes of *B. duncani*, 2,645 (63% of the total) were multi-exon and 3,200 had a functional annotation. The *B. duncani* nuclear genome expresses two 18S rRNA encoding genes, one on Chromosome I (coordinates: 932670 to 934438) and the second on Chromosome III (coordinates 445698 to 447466), three 5S rRNAs genes in tandem on Chromosome II (coordinates: 805406 to 805517; 815064 to 815175 and 824722 to 824833) as well as 46 tRNAs distributed on all five chromosomes and six others in unplaced contigs. Transcript analysis using BUSCO v5 (23) showed the transcripts’ annotations matching with 88.6% of the transcripts in the apicomplexa_odb10 database (80.5% single-copy, and 8.1% duplicated), whereas 5.8% are fragmented and 5.6% are missing. BUSCO analysis further showed that the *B. duncani* protein annotations match 87.7% of the proteins in the apicomplexa_odb10 database (79.6% single-copy, and 8.1% duplicated), whereas 5.6% are fragmented and 6.7% are missing. Only 1.2% of the genome is considered repetitive by RepeatMasker. Repeats were detected in the following categories: simple repeats (0.79%), low complexity repeats (0.09%), small RNA (0.29%) and other interspersed repeats (0.03%).

### Genome-wide comparison of *B*. *duncani* gene models with those of other Apicomplexa

The *B. duncani* nuclear genome has a GC content of 37.32%, which is similar to that of other piroplasms (25) and two times higher than that of *P. falciparum* (26). This property makes *B. duncani* an attractive model system to study gene function in heterologous systems using standard molecular biology and genetic tools as well as in the parasite itself to examine the importance of these genes involved in parasite survival, virulence and susceptibility to antiparasitic drugs (11). Main genome and gene statistics of *B. duncani* and other Apicomplexa are shown in **Table 2**. Observe that the genome size, the number of chromosomes, and the number of protein coding genes in *B. duncani* is higher than those in *B. microti, B. bovis, T. parva, C. parvum* and *T. annulate,* which could be also attributed to a more complete genome assembly for *B. duncani.* An approximate 60% of the *B. duncani* genome encodes proteins, the percentage of genes with introns is ∼63%, and the average number of exons per gene is 3.1.

Out of the total *B. duncani* genes, 3,484 mapped to chromosomes while the rest were located on unplaced contigs. Genes were evenly distributed across the five chromosomes with gene poor regions localized to telomeric ends most notably on one side of chromosome III (Fig. 3A, black bars). Mapping of *B. duncani* genes to orthology groups in OrthoMCL revealed 547 (14% of chromosomal genes) genes that are unique to *B. duncani*, out of these 198 included genes with at least one paralog suggesting possible expanded families unique to this species (Fig 3A, red and orange bars). Additionally, 1,733 genes exhibited no orthology outside the piroplasmida and/or haemosporidia including 844 genes that were conserved between *B. duncani* and all other piroplasmida, 118 that had at least one ortholog in another piroplasmid and 811 that had at least one ortholog between *B. duncani* at least one piroplasmida or haemosporidia (Fig. 3A, green, blue and purple bars). Comparison of orthology groups across different *Babesia* species revealed a core set of 1,943 orthology groups (Fig. 3B). Interestingly, *B. microti* contained the largest number of unique orthology groups and observation that is also supported by its phylogenetic relationship to other *Babesia* (See below). Not surprisingly the core set of orthology groups was less (1, 107) when comparing *B. duncani* to other apicomplexa (Fig. 3C). The largest number of shared orthology groups (510) was between *B. duncani*, *T. parva* and *P. falciparum*. Genome size and gene content comparing *B. duncani* with other Apicomplexa is summarized in Fig. 3D. Gene content did not reveal any surprises and largely correlated with genome size. A phylogenetic analysis of the same organisms shown in Fig. 3D revealed that *B. duncani* forms a well-supported clade with *B. bovis* and *B. bigemina* while *B. microti* diverged early compared to other piroplasmida include *T. parva* and *T. annulate*.

**Figure 3.**
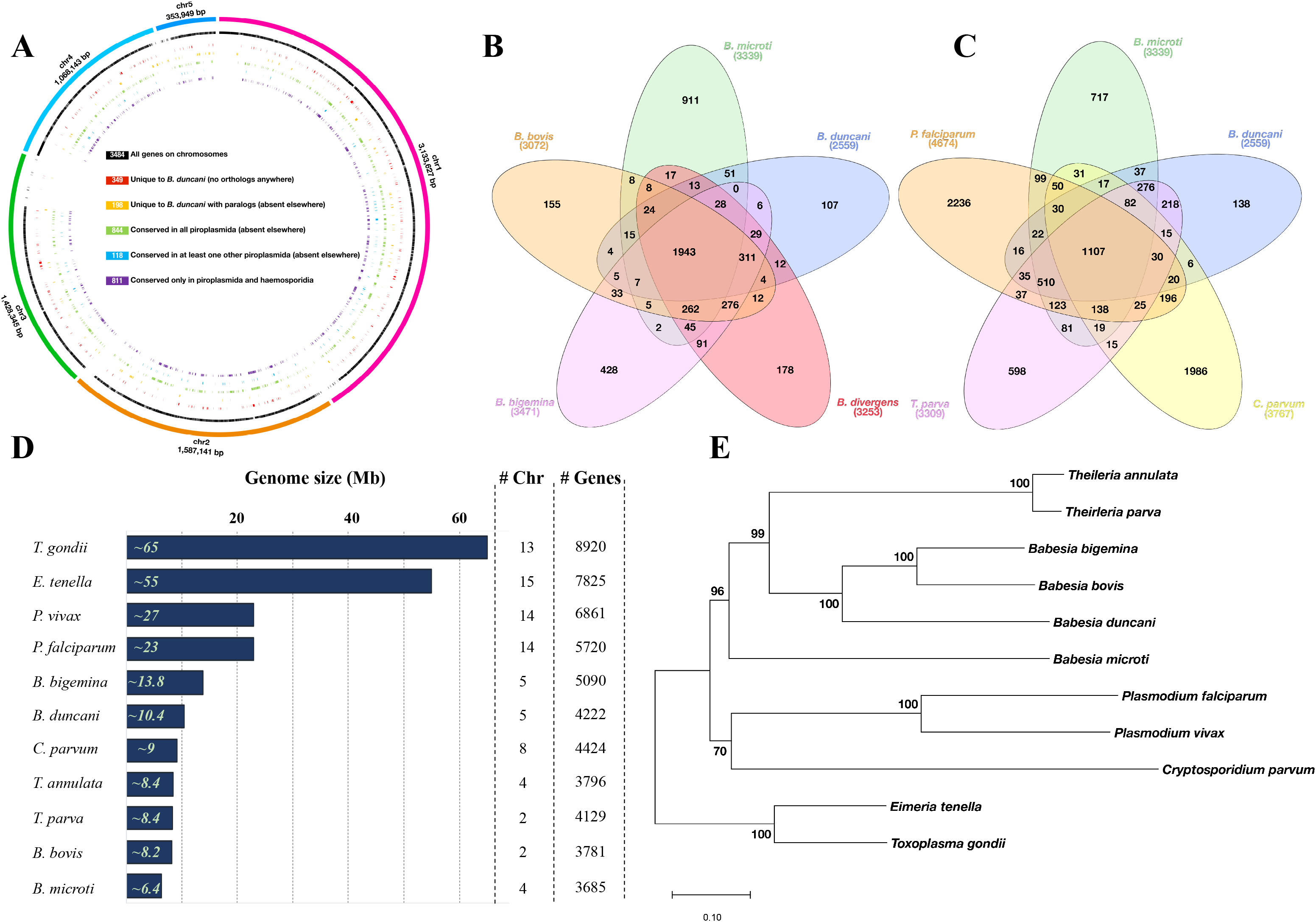
**A.** Circos plot depicting *B. duncani* gene distribution across the five chromosomes. Outer circle depicts the chromosomes, in black all genes, in red all genes unique to *B. duncani*, in orange unique genes that have paralogs, in green genes that are conserved in piroplasmida and do not have orthologs elsewhere, in blue genes with some conservation in piroplasmida and absent elsewhere and in purple are genes that are conserved in both piroplasmida and haemosporidia but absent elsewhere. **B.** Venn diagram showing the overlap between orthology groups from *Babesia sp.* (*B. duncani* in blue, *B. divergens* in red, *B. bigemina* in pink, *B. bovis* in orange, *B. microti* in green). **C.** Venn diagram showing the overlap between orthology groups from *B. duncani* in blue, *B. microti* in green, *Cryptosporidium parvum* in yellow, *Theileria parva* in pink and *Plasmodium falciparum* in orange,). **D.** schematic showing genome sizes in megabases of various Apicomplexa. Gene numbers include both coding and non-coding genes. **E.** Phylogenetic tree of selected Apicomplexans.

Syntenic genome analyses were conducted between *B. duncani* and four other hemoparasites, *B. microti, T. parva, B. bovis* and *B. bigemina* and illustrated as Circos plots in which blue arcs indicate syntenic regions and red arcs indicate inverted syntenic regions. The intensity of the red/blue color is proportional to the amount of collinearity. As shown in Fig. 4, *B. microti* is the least syntenic with *B. duncani* compared to the other three species, consistent with our phylogenetic tree analyses. The comparative analysis between *B. duncani* and *T. parva* indicates, for instance, a ∼1Mb syntenic inversion between the respective chromosomes 1, and a ∼0.5Mb collinear region between chromosome 2 of *B. duncani* and chromosome 4 of *T. parva.* The comparison between *B. duncani* and *B. bovis* indicates that almost the entire chromosome 2 of *B. duncani* is syntenic to the second half of chromosome 3 in *B. bovis*. Finally, the comparative analysis between *B. duncani* and *B. bigemina* indicates strong collinearity of chromosome 1 in *B. duncani* and chromosome 2 in *B. bigemina,* as well as a large synteny between chromosome 4 in *B. duncani* and chromosome 4 in *B. bigemina*.

**Figure 4.**
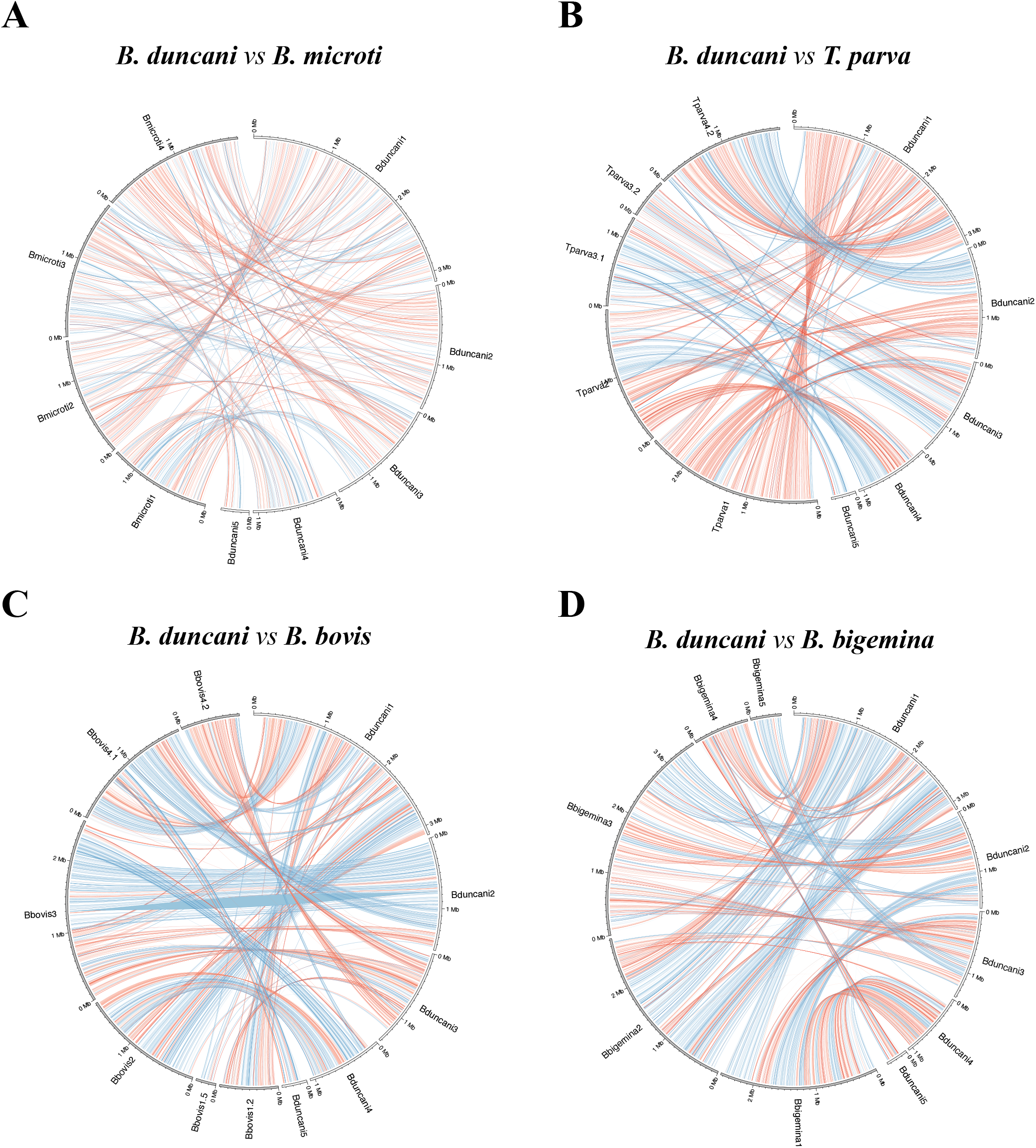
Circos synteny plots. The chromosomes of *B. duncani* are illustrated on the right semicircle on all circular plots, and the chromosomes of the other organisms are on the left semicircle (**A**: *B. microti,* **B**: *T. parva,* **C**: *B. bovis,* **D**: *B. bigemina*); blue arcs indicate syntenies, red arcs indicate syntenies involved in a reversals; the intensity of the color is proportional to the level of collinearity; the number after the species’ name refers to the chromosome number (when chromosomes are broken into pieces, fragments are numbered); chromosomes that had no significant syntenies were not included.

The metabolic atlas of *B. duncani* was reconstituted from the annotated genome and transcriptomic profile with most housekeeping functions sharing homologs in other Babesia species, *Theileria* and *Plasmodium*. All enzymes of the glycolytic pathway and the tricarboxylic acid cycle were found in the annotated proteome (Tables 3 and 4). Transcriptomic analyses suggest that *B. duncani* operates a standard glycolytic cycle whereas its TCA cycle might follow a reductive process similar to that previously reported for *B. microti* (25). Whereas eight of the ten genes encoding enzymes in the glycolytic pathway are expressed at high levels, the enolase and lactate dehydrogenase genes are expressed at low levels (Table 3). On the other hand, two critical enzymes of the *B. duncani* TCA cycle, succinyl dehydrogenase and succinyl-CoA synthetase alpha subunit, were found to be not expressed during the intraerythrocytic life cycle of the parasite. These findings suggest that the TCA cycle of *B. duncani* might work in a reductive way from alpha-ketoglutarate generated from glutamine. The pathway from succinate to oxaloactetate is connected to the ubiquinone pool and mitochondrial ATPase activity through succinate dehydrogenase and malate:quinone oxidoreductase (BdWA1_002484). Unlike the *B. microti* Lactate dehydrogenase (BmLDH), which has been shown to be of host origin (25), the lactate dehydrogenase (LDH) of *B. duncani* is similar to LDH enzymes of most apicomplexan parasites whose evolution has been suggested to derive from a malate dehydrogenase (27). The *B. duncani* LDH is thus likely to function primarily in the production of lactate.

**Table 3.** Predicted enzymes of the glycolytic pathway of *B. duncani*.

| Glycolysis Steps | Enzyme | Gene ID | Protein Length | RNA expression level (TPM) |
| --- | --- | --- | --- | --- |
| Glucose |  |  |  |  |
| ↓ | ← Hexokinase | BdWA1_000031-T1 | 486 | 73 |
| Glucose-6P |  |  |  |  |
| ↓ | ← Phosphoglucose Isomerase | BdWA1_002840-T1 | 422 | 511 |
| Fructose-6P |  |  |  |  |
| ↓ | ← 6-Phosphofructokinase | BdWA1_003208-T1 | 1313 | 506 |
| Fructose-1,6P2 |  |  |  |  |
| ↓ | ← Fructose-1,6-bisphosphate aldolase | BdWA1_000083-T1 | 356 | 1884 |
| Glyceraldehyde-3P |  |  |  |  |
| ↓ | ← Glyceraldehyde-3P dehydrogenase | BdWA1_001300-T1 | 336 | 8497 |
| Glycerate-1,3P2 |  |  |  |  |
| ↓ | ← Phosphoglycerate Kinase | BdWA1_002139-T1 | 416 | 2018 |
| Glycerate-3P |  |  |  |  |
| ↓ | ← Phosphoglycerate mutase | BdWA1_003538-T1# | 299 | 50 |
| Glycerate-2P |  |  |  |  |
| ↓ | ← Enolase | BdWA1_003622-T1 | 442 | 15 |
| Phosphoenolpyruvate |  |  |  |  |
| ↓ | ← Phosphoenolpyruvate carboxykinase | BdWA1_000439-T1 | 556 | 425 |
| Pyruvate |  |  |  |  |
| ↓ | ← Lactate dehydrogenase | BdWA1_000746-T1 | 320 | 22 |
| Lactate |  |  |  |  |

#: BdWA1\_001702-T1

**Table 4.** Predicted enzymes of the TCA cycle of *B. duncani*.

| Krebs Cycle steps | Enzyme | Predicted Gene ID | Protein Length | C/M/S | TM | RNA expression level (TPM) |
| --- | --- | --- | --- | --- | --- | --- |
| Citrate |  |  |  |  |  |  |
| ↓ | ← Aconitate Hydratase | BdWA1_001117-T1 | 806 | M | - | 745 |
| Isocitrate |  |  |  |  |  |  |
| ↓ | ← Isocitrate dehydrogenase | BdWA1_001763-T1 | 454 | M | - | 773 |
| 2-oxaloglutarate |  |  |  |  |  |  |
| ↓ | ← 2-Oxoglutarate dehydrogenase | BdWA1_000507-T1 | 940 | M | - | 293 |
| Succinyl-CoA |  |  |  |  |  |  |
| ↓ | ← Succinyl-CoA synthetase | BdWA1_001709-T1#<br>BdWA1_002351-T1 | 309<br>423 | M<br>M | -<br>- | 0.58<br>204 |
| Succinate |  |  |  |  |  |  |
| ↓ | ← Succinate dehydrogenase | BdWA1_004214-T1* ($\alpha$)<br>BdWA1_000189-T1 ($\beta$) | 617<br>280 | M<br>- | -<br>- | 0<br>93 |
| Fumarate |  |  |  |  |  |  |
| ↓ | ← Fumarase | BdWA1_000547-T1 | 578 | - | - | 175 |
| Malate |  |  |  |  |  |  |
| ↓ | ← Malate dehydrogenase | BdWA1_002484-T1 | 489 | - | 1 | 165 |
| Oxaloacetate |  |  |  |  |  |  |
| ↓ | ← Citrate synthase | BdWA1_003132-T1 | 553 | - | - | 485 |

\* Identical proteins: BdWA1\_004074-T1, BdWA1\_003754-T1, BdWA1\_003433-T1

Of the 4,170 proteins encoded by the *B. duncani* nuclear genome, 479 (11.4% of CDSs) harbor an N-terminal signal peptide thus highlighting the importance of protein secretion in this parasite’s intraerythrocytic life cycle. Our analysis further identified 106 proteins (2.5% of CDSs) encoded by the *B. duncani* nuclear genome and targeted to the mitochondria. 781 proteins (19% of CDSs) have at least one transmembrane domain with five proteins predicted to have 18 transmembrane domains each (BdWA1_000001, BdWA1_000770, BdWA1_002261, BdWA1_003351 and BdWA1_003495). Unlike *P. falciparum*, which expresses 13 aspartic proteases, four of which (Plasmepsins I-IV) are involved in various steps of hemoglobin degradation in the digestive vacuole (28), *B. duncani* encodes seven aspartic proteases of which only four are expressed during the intraerythrocytic life cycle of the parasite as determined by RNAseq. Of these, BdWA1_003023 shares 50% identity with DdiI (DNA damage-inducible protease 1), BdWA1_003366 shares ∼45% identity with plasmepsins IX and X involved in invasion and/or egress, BdWA1_002231 showed the highest sequence similarity with plasmepsins V involved in protein transport and dense granule protein maturation, and BdWA1_001984 shares homology with all *P. falciparum* aspartic proteases in the range of 23 to 27% identity.

The most conserved genes between *B. duncani* and other *Babesia* species are those involved in DNA replication, transcription, and protein translation. Our analysis identified 19 members of the ApiAP2 family that belong to different orthology groups. RNA-seq data show that 15 of the 19 *BdAP2* genes are expressed above background during the intraerythrocytic cycle of the parasite (Table 5), suggesting that the four remaining BdAP2 genes may be important for the development of the parasite in the tick vector. The annotation of the *B. duncani* proteome further identified 22 general factors involved in the recruitment of RNA polymerases I, II and III. The translation machinery of *B. duncani* includes 25 aminoacyl-tRNA synthetases and 187 ribosomal proteins, 87.2% of which are expressed (i.e., TPM > 10.0) during the intraerythrocytic life cycle of the parasite in human red blood cells.

**Table 5.** Predicted GPI-anchored proteins of *B. duncani*.

|  |  |  |  | SignalP 5.0 | PredGPI |  | BLAST |  |
| --- | --- | --- | --- | --- | --- | --- | --- | --- |
| GPI-AP ID | Protein ID | Length (aa) | RNA expression (TPM) | Score | Prediction | Specificity | Description | Taxonomy distribution |
| BdGPI01 | BdWA1_000266-T1 | 498 | 22 | 0.604043 | Highly probable | 100% | AAY83299, putative novel antigen from Babesia sp. WA1 | WA1 |
| BdGPI02 | BdWA1_000268-T1 | 514 | 40 | 0.604043 | Highly probable | 100% | AAY83299, putative novel antigen from Babesia sp. WA1 | WA1 |
| BdGPI03 | BdWA1_000269-T1 | 506 | 1.8 | 0.604043 | Highly probable | 100% | AAY83299*, putative novel antigen from Babesia sp. WA1 | WA1 |
| BdGPI04 | BdWA1_000270-T1 | 514 | 34 | 0.604043 | Highly probable | 100% | AAY83299, putative novel antigen from Babesia sp. WA1 | WA1 |
| BdGPI05 | BdWA1_000500-T1 | 189 | 24 | 0.948747 | Highly probable | 99.9% | No significant similarity found | NA |
| BdGPI06 | BdWA1_000716-T1 | 236 | 1.4 | 0.989215 | Probable | 99.8% | BmGPI14, B. microti R1 | Piroplasmida |
| BdGPI07 | BdWA1_001123-T1 | 377 | 278 | 0.846581 | Highly probable | 100% | AAY83302*, putative novel antigen from Babesia sp. WA1 | WA1 |
| BdGPI08 | BdWA1_001649-T1 | 457 | 123 | 0.904527 | Weakly probable | 99.4% | hypothetical protein | Piroplasmida (not Bmicroti) |
| BdGPI09 | BdWA1_001913-T1 | 729 | 285 | 0.992360 | Highly probable | 100% | carbonic anhydrase | Apicomplexa (+) |
| BdGPI10 | BdWA1_001963-T1 | 1851 | 5 | 0.972058 | Probable | 99.6% | hypothetical protein | WA1 |
| BdGPI11 | BdWA1_002110-T1 | 330 | 32 | 0.998113 | Highly probable | 100% | No significant similarity found | NA |
| BdGPI12 | BdWA1_002112-T1 | 378 | 160 | 0.980121 | Highly probable | 99.9% | No significant similarity found | NA |
| BdGPI13 | BdWA1_002223-T1 | 242 | 1656 | 0.962141 | Highly probable | 99.9% | AAY83296*, putative novel antigen from Babesia sp. WA1 | WA1 |
| BdGPI14 | BdWA1_002224-T1 | 518 | 408 | 0.969359 | Highly probable | 99.9% | AAY83296, putative novel antigen from Babesia sp. WA1 | WA1 |
| BdGPI15 | BdWA1_002538-T1 | 631 | 17 | 0.985101 | Probable | 99.8% | No significant similarity found | NA |
| BdGPI16 | BdWA1_003372-T1 | 122 | 8441 | 0.736131 | Highly probable | 99.9% | No significant similarity found | NA |
| BdGPI17 | BdWA1_003373-T1 | 127 | 8200 | 0.758117 | Highly probable | 99.9% | No significant similarity found | NA |

\* indicates proteins with more than 95% identity with BLASTP. (+) for proteins with more similarities outside the main taxonomic group with less coverage and lower score. Color code highlight paralogues from the same group.

**Table 6.**
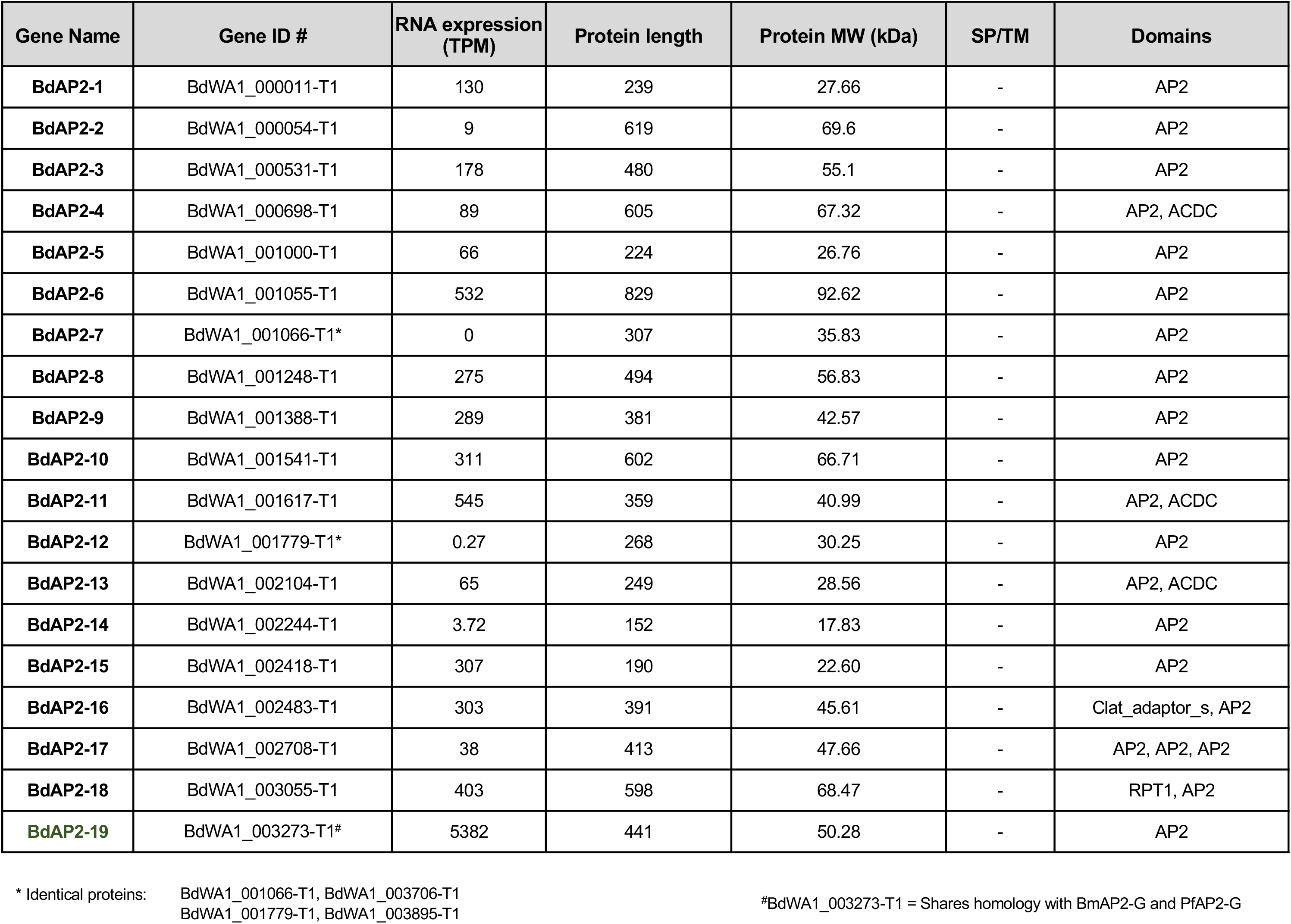
Predicted AP2 transcription factors of *B. duncani*.

| Gene Name | Gene ID # | RNA expression (TPM) | Protein length | Protein MW (kDa) | SP/TM | Domains |
| --- | --- | --- | --- | --- | --- | --- |
| **BdAP2-1** | BdWA1_000011-T1 | 130 | 239 | 27.66 | - | AP2 |
| **BdAP2-2** | BdWA1_000054-T1 | 9 | 619 | 69.6 | - | AP2 |
| **BdAP2-3** | BdWA1_000531-T1 | 178 | 480 | 55.1 | - | AP2 |
| **BdAP2-4** | BdWA1_000698-T1 | 89 | 605 | 67.32 | - | AP2, ACDC |
| **BdAP2-5** | BdWA1_001000-T1 | 66 | 224 | 26.76 | - | AP2 |
| **BdAP2-6** | BdWA1_001055-T1 | 532 | 829 | 92.62 | - | AP2 |
| **BdAP2-7** | BdWA1_001066-T1* | 0 | 307 | 35.83 | - | AP2 |
| **BdAP2-8** | BdWA1_001248-T1 | 275 | 494 | 56.83 | - | AP2 |
| **BdAP2-9** | BdWA1_001388-T1 | 289 | 381 | 42.57 | - | AP2 |
| **BdAP2-10** | BdWA1_001541-T1 | 311 | 602 | 66.71 | - | AP2 |
| **BdAP2-11** | BdWA1_001617-T1 | 545 | 359 | 40.99 | - | AP2, ACDC |
| **BdAP2-12** | BdWA1_001779-T1* | 0.27 | 268 | 30.25 | - | AP2 |
| **BdAP2-13** | BdWA1_002104-T1 | 65 | 249 | 28.56 | - | AP2, ACDC |
| **BdAP2-14** | BdWA1_002244-T1 | 3.72 | 152 | 17.83 | - | AP2 |
| **BdAP2-15** | BdWA1_002418-T1 | 307 | 190 | 22.60 | - | AP2 |
| **BdAP2-16** | BdWA1_002483-T1 | 303 | 391 | 45.61 | - | Clat_adaptor_s, AP2 |
| **BdAP2-17** | BdWA1_002708-T1 | 38 | 413 | 47.66 | - | AP2, AP2, AP2 |
| **BdAP2-18** | BdWA1_003055-T1 | 403 | 598 | 68.47 | - | RPT1, AP2 |
| **BdAP2-19** | BdWA1_003273-T1<sup>#</sup> | 5382 | 441 | 50.28 | - | AP2 |

\* Identical proteins: BdWA1\_001066-T1, BdWA1\_003706-T1 BdWA1\_001779-T1, BdWA1\_003895-T1
<sup>#</sup>BdWA1\_003273-T1 = Shares homology with BmAP2-G and PfAP2-G

### New multigene families of *B. duncani*

The extremely high virulence of *B. duncani* and the cytokine storm it triggers in immunocompetent mice and hamsters suggest that the parasite produces virulence factors during its blood stage development that are delivered into the host and trigger a strong host response (29). Interestingly, our analysis of the *B. duncani* predicted genes identified 747 genes (18% of the protein coding genes) that belong to multigene families (MGF) each with at least three gene members. These families can be divided into two classes: BdUMGF (<u>u</u>nique to *B. duncani*) and BdOMGFs (with <u>o</u>rthologs in other *Apicomplexa*). The BdUMGF class includes 327 genes grouped into 52 gene families. The largest family in this group (BdUMGF1) consists of 51 members (Fig. 5A and 5B). The BdOMGFs comprises 420 genes grouped into 105 genes families each belonging to an orthologous group among *Apicomplexa*. The largest family, BdOMGF1, belongs to orthology group OG6_101304 that includes 288 orthologs in *Theileria spp*, majority of which are hypothetical proteins, 5 Tpr-related proteins in *B. microti*, 5 hypothetical proteins in *B. bovis* and 3 members of the bir1, PIR and YIR in *P. yoelii.* The components of each of the gene families and their properties are available in the supplemental datasets. The chromosomal location of the genes in the three largest BdUMGF and BdOMGF families, as well as the GPI and AP families, is illustrated in Figure 5C. Observe that (i) genes in the BdUMGF1 family (indicated by the red color) are strongly colocalized and on the same strand at the telomeres of chromosome 1 and chromosome 4, which also appear to be in close proximity in the 3D model (Figure 2D); (ii) all genes in the BdUMGF3 family (indicated by the dark blue color) are clustered together on the same strand in the middle of chromosome 3; (iii) all genes in BdOMGF2 family (indicated by the magenta color) are clustered together on the same strand in the middle of chromosome 1; (iv) genes in the GPI and AP family are uniformly distributed along the chromosomes. Interestingly, genes in the BdUMGF family are not only in closed proximity to each other at the chromatin 3D level, but also seem to be under transcriptional repression as a result of their chromosomal location (Fig 5A and 5B). This may indicate the presence of a possible heterochromatin cluster surrounding these genes similar to what has been described for the *var* or *SICAvar* gene families of *P. falciparum* and *P. knowlesi,* respectively. The *P. falciparum* Var gene family consists of ∼7x genes that encode a major virulence antigen, PfEMP1, with a single variant expressed in each infected red blood cell. PfEMP1-mediated antigenic variation has been suggested to play a role in *P. falciparum* immune evasion. The control of mutually exclusive var gene expression in the parasite relies on epigenetic and chromatin structure changes that are critical for pathogenesis and immune evasion. Considering the similarity observed between the var and BdUMGF gene families at the chromatin 3D and gene expression level, it is likely that genes that belong to BdUMGF are involved in antigenic variation, explaining to some extend the high virulent status of *B. duncani*.

**Figure 5.**
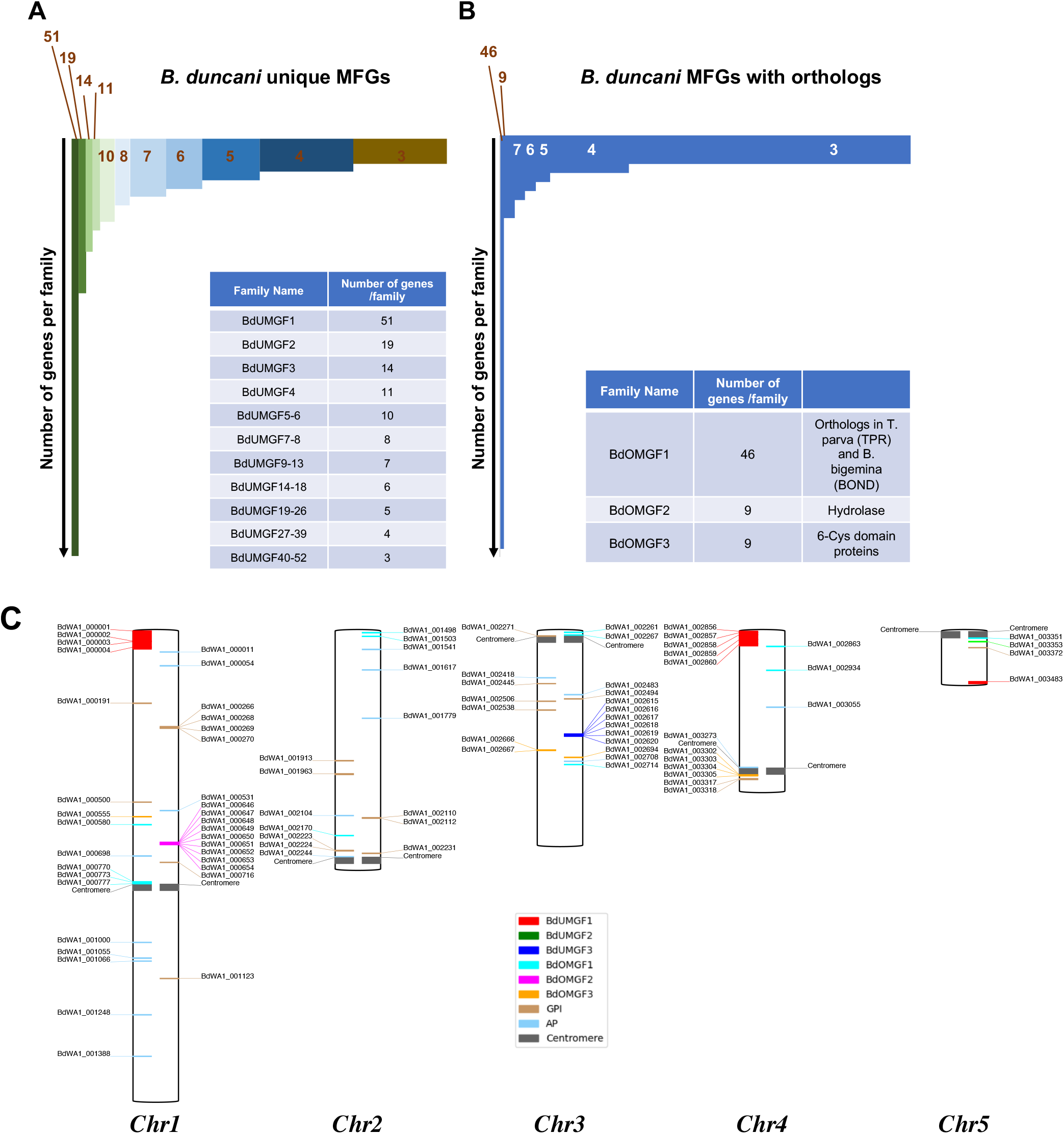
Multigene families and chromosomal localization in *B. duncani*. **A.** gene families unique to *B. duncani*. **B:** gene families with orthologs in other Apicomplexa. **C:** Localization of the genes in the gene families BdUMGF1, BdUMGF2, BdUMGF3, BdOMGF1, BdOMGF2, BdOMGF3, GPI, AP on the five *B. duncani* chromosomes (genes localized on unplaced contigs are ignored); genes on the right side of a chromosome are on the positive strand, genes on the left side are on the negative strand.

### GPI-anchored proteome of *B. duncani*

Studies in apicomplexan parasites identified several GPI-anchored proteins (GPI-AP) that function as major antigens. These include MSP proteins in *P. falciparum*, Bd37 in *B. divergens* or BmSA1/BmGPI12 in *B. microti* (30–33). Interestingly, none of these proteins has orthologues in the *B. duncani* proteome. In silico prediction of the GPI-proteome of *B. duncani* identified 17 GPI-anchored proteins, the majority of which are unique to the parasite and 14 of which are expressed during the intraerythrocytic life cycle (TPM >10) (Table 5). Seven of the BdGPI-AP proteins share sequence similarities with previously described putative *B. duncani* antigens (Table 5). The *B. duncani* BdGPI06 (BdWA1_000716) is the only GPI-AP of this parasite with an ortholog in *B. microti* (BmGPI4) and is also conserved among other piroplasmida. Orthologs of BdGPI08 (BdWA1_001649-T1) are found in other piroplasmida but not *B. microti*. Interestingly, among the *B. duncani* GPI-anchored proteins identified in our analysis, the BdGPI09 protein shares homology with members of the carbonic anhydrase (CA) superfamily. These metalloenzymes play a crucial physiological function by catalyzing the reversible reaction of CO_2_ hydration to HCO_3_^−^ and H^+^ (*CO*+2*HO*2⇋*HCO*+3−*H*+), which is essential for pH homeostasis, secretion of electrolytes and various biosynthetic processes. While the finding of a CA enzyme as a GPI-anchored protein is unique among *Apicomplexa*, there is a precedent for this in mammals including the human and mouse CA IV (34). Thus, BdGPI09 could play a critical role in the regulation of pH cytoplasm in *B. duncani* and could represent an excellent target for the development of antiparasitic drugs. Another unique feature of the GPI-anchored proteins of *B. duncani* is that several of their encoding genes are found to be duplicated locally in the *B. duncani* genome with six of these proteins being members of the BdUMGF9 specific family (Fig. 5).

### Regulation of gene expression in *B. duncani*

Our RNA-seq analysis, which was used to refine the genome annotation, helped determine the expression levels of encoded gene to gain insights into their importance during the parasite life cycle in mammalian blood or the tick vector families as well as to provide evidence for function metabolic pathways that could be targeted for the development of novel therapeutic strategies. Using transcript abundance values based on the mean depth of coverage (see *Experimental procedures*), we found that the total range of transcriptional activity captured using the continuous *in vitro* growth conditions varied by more than four orders of magnitude (Fig. 6 and **supplemental datasets**). Overall, RNA-Seq data captured close to 90% of the predicted annotated genes in the assembled *B. duncani* genome (Fig. 6A and B), indicating that most genes in the *B. duncani* genome are needed for parasite survival in host red blood cells. Among the most highly expressed genes were genes involved in translation, ubiquitin proteasome system, cell cycle, ATP hydrolysis-coupled proton transport, carbohydrate metabolic process as well as histone core proteins indicating active metabolic activity and maintenance of the parasite by standard housekeeping genes (Fig. 4D). Amongst the 422 genes that were found to be silenced during the intraerythrocytic life cycle in vitro, there were genes involved in microtubule-based processes that are known to be up-regulated during sexual differentiation in other apicomplexan parasites (35). Several kinases, phosphatases, mRNA binding proteins as well as a few transcription factors including 3 AP2 genes, know to regulate transcriptional initiation in other apicomplexan parasite, were also found in this group of genes that were not expressed and could have a role in sexual differentiation or life cycle progression in the tick vector. We then exploited this dataset further to mine and identify additional molecular components that could be critical to the survival of the parasite. Of the reads that mapped uniquely against the babesia genome, ∼83.91% mapped to predicted protein-coding genes and 23.6% mapped within 300-bp up- or downstream of genes as well as within intergenic regions, demarcating possible UTRs and lncRNAs respectively. Long non-coding RNAs (lncRNAs) are non-protein coding transcripts that are generally regulated and processed similarly to mRNAs and have been shown to play a critical role in biology including cell differentiation throughout changes in epigenetics and chromatin structures. Their roles have also been implicated in the regulation of genes involved in antigenic variation in human malaria parasites (36).

**Figure 6.**
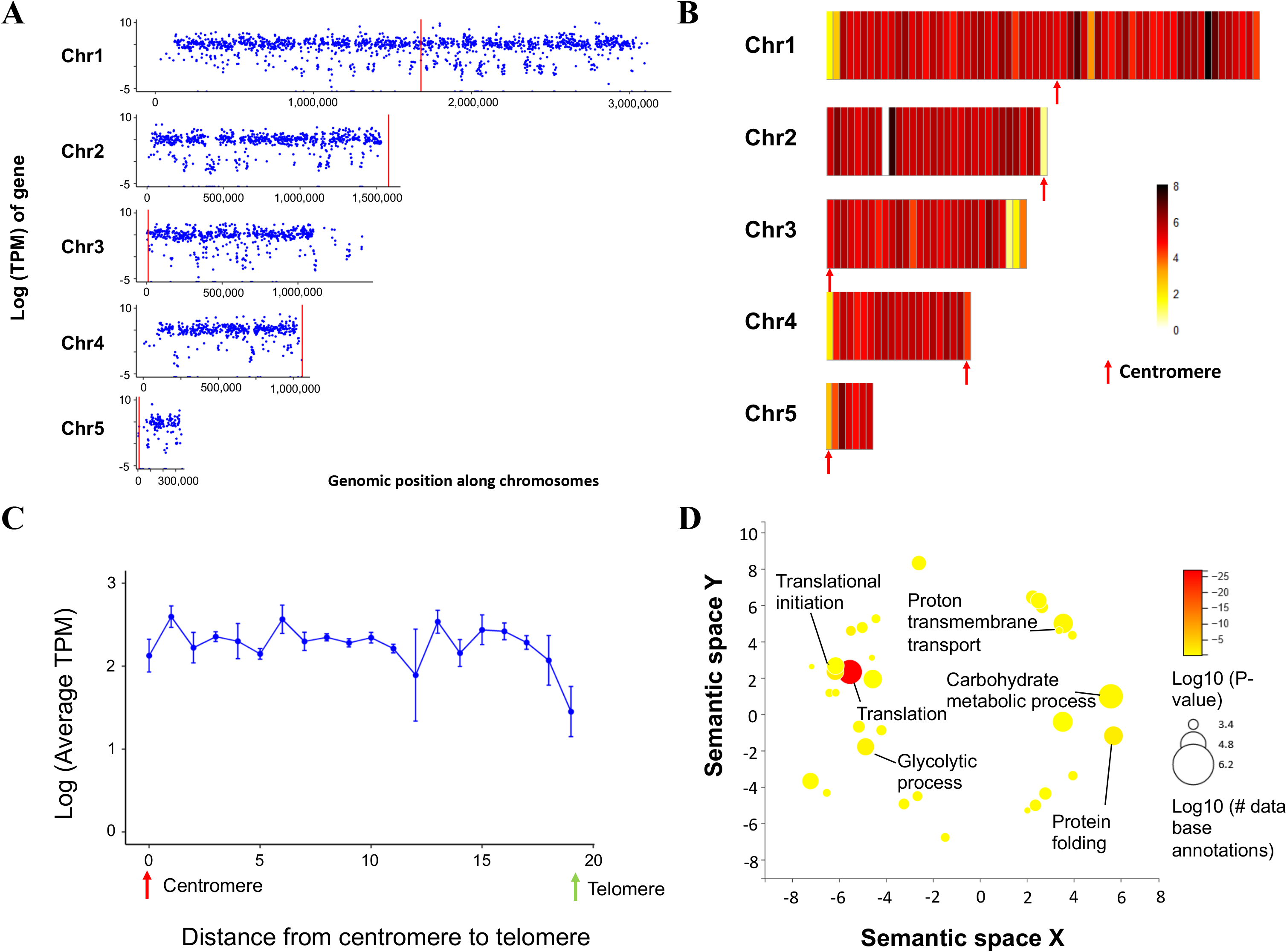
RNA-seq data in *B. duncani*. **A.** Logarithms of the TPM counts were used as expression value for each genes across the 5 chromosomes using the R package ggplot2 **B.** RNA-seq data as Normalized heatmaps across the 5 chromosomes. Chromosomes were divided into bins at 50-kb bins and the average of the log TPM of genes within each bin was calculated. **C.** Relation between gene expression and distance from the centromeres to the telomeres. Chromosomes were divided into 20 bins. For each bin, the average gene expression value was plotted. Error bars denote the range of expression values within each bin for each chromosome. **D.** Gene ontology enrichment of the most highly expressed genes. GO was calculated using the R package TopGO with the weight01 algorithm and the biological process tree. REVIGO was used to visualize the GO results of the most highly expressed genes (500TPM or higher).

We next examined the possible relationship between gene expression and genome organization. We therefore binned the genes of *B. duncani* into 20 groups based on their distance from the centroid of telomeres and calculated the average gene expression using our RNA-seq data and plotted these values against the average distances from the telomere centroid (Fig. 6C). We also color coded the bins on normalized average gene expression values across the whole chromosomes (Fig 6B). Similar to what we observed in *the P. falciparum* and *P. knowlesi* genomes, two apicomplexan parasites that possess gene families involved in antigenic variation, we detected a significant relationship between gene expression and 3D location relative to the telomeres. A significant decrease in the expression of the genes localized near the centromeres was also observed. Gene repression was noticeable near one of the telomeric ends of chromosome 1, 3, 4 and 5, which harbor clusters of genes belonging to the *B. duncani* BdUMGF1 multigene family (Fig. 5C). None of these features was detected in the *B. microti* genome. As previously noted, this result suggests that the *B. duncani* genome may contain an heterochromatin cluster near the telomere ends allowing for mono-allelic expression of the BdUMGF1.

### Antifolates are potent inhibitors of *B. duncani* intraerythrocytic development

The recent development of a continuous *in vitro* culture system for *B. duncani* in human red blood cells made it possible to investigate the susceptibility of this parasite to recommended babesiosis therapies (11). These studies revealed that the parasite was inherently less susceptible to atovaquone (IC_50_ ∼0.5 µM), azithromycin (IC_50_ ∼5 µM), clindamycin (IC_50_ ∼20 µM) and quinine (IC_50_ ∼12 µM) than *P. falciparum* (37, 38). Mining of the annotated *B. duncani* proteome identified several potential drug targets (Fig. 7). One of these, is the dihydrofolate reductase-thymidylate synthase (DHFR-TS) Fig. 8A. Residue T71 in *B. duncani* DHFR-TS (BdDHFR-TS) sequence is equivalent to residue 108 in DHFR-TS from *P. falciparum* (PfDHFR-TS). Sensitivity or elevated resistance to pyrimethamine in *P. falciparum* is associated in serine or asparagine substitutions in position 108 at PfDHFR-TS sequence, receptivity (39) (Fig. 8B). Consistent with our genome analysis, drug sensitivity and *in vitro* safety assays revealed that *B. duncani* is sensitive to pyrimethamine and WR-99210 with IC_50_ values of 940 and 0.58 nM, respectively, with both compounds showing excellent *in vitro* therapeutic indices when tested against four human cell lines (**Fig. 9C-D**). The specificity of inhibition by these drugs was further examined biochemically using recombinant DHFR-TS enzymes from *B. duncani* and *B. microti* (BmDHFR-TS) (**Fig. 9E-F**). WR-99210 was found to inhibit BdDHFR-TS and BmDHFR-TS enzymatic activity with IC_50_ of 3.9 and 6.8 nM, respectively. Our biochemical assays results showed that both enzymes have similar kinetic properties (*K_m_* and V_max_), but different turnover number (k_cat_). This variation led to improved catalytic efficiency (k_cat_/*K_m_*) on both substrates (DHF and NADPH) of BmDHFR-TS over BdDHFR-TS (**Fig. 9F**). All together these findings support the use of antifolates as potential therapies for the treatment of human babesiosis.

**Figure 7.**
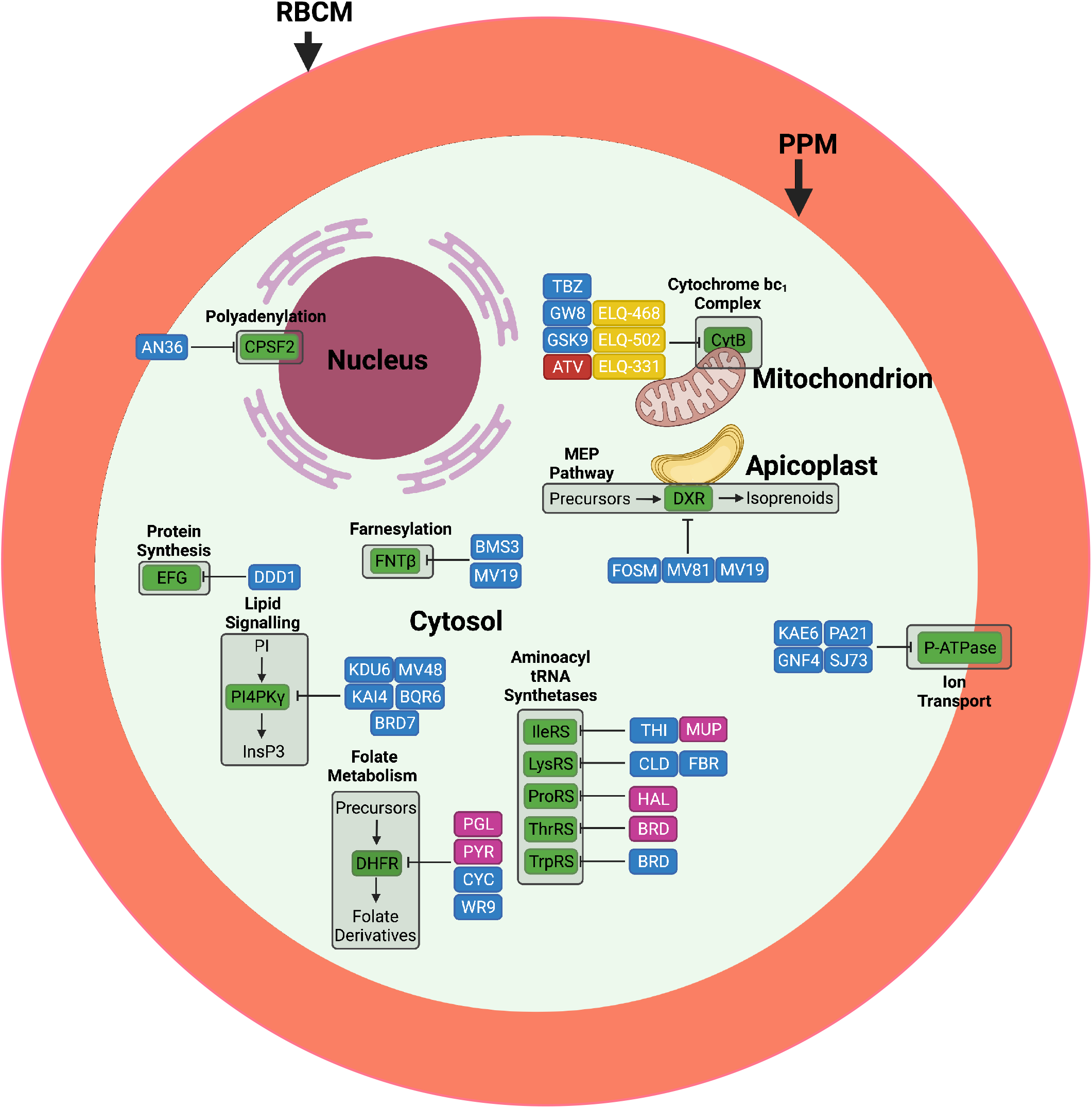
Putative molecular targets of *B. duncani* and candidate. Drugs are categorized as follows: (Red) Effective against Babesia parasites and used clinically; (Yellow) Effective against *Babesia* parasites but have not been evaluated clinically; (Pink) Clinically approved drugs for other diseases but have not been tested against Babesia parasites; and (Blue) Drug under clinical evaluation for the treatment of other diseases but have not yet been tested against *Babesia* parasites inhibitors. Drug and protein abbreviations are as follows: Isoleucyl tRNA Synthase (Iso-TRNAS), Lysyl tRNA Synthase (Lys-TRNAS), Prolyl tRNA Synthase (Pro-TRNAS), Theronyl/alanyl tRNA Synthase (Thre-TRNAS), Tryptophanyl tRNA Synthase (Tryp-TRNAS), Dihydrofolate reductase (DHFR), P-ATPase (P-type ATPase), CytB (Cytochrome b-c1 complex subunit 7 superfamily), Translation EFG/EF2, Elongation factor EFG (EFG), Phosphatidylinositol 4-kinase gamma (PI4PKgamma), Farnesyltransferase subunit beta (FNTB), Polyadenylation specificity factor subunit 2 (CPSF2), MEP Synthase (DXR), Febrifugine (FBR), Halofuginone (HAL), Cladosporin (CLD), Mupirocin (MUP), Borrelidin (BRD), Indolmycin (IND), DDD107498 (DDD1), KDU691 (KDU6), MMV048 (MV48), KAI407 (KAI4), BQR695 (BQR6), BRD73842 (BRD), Atovaquone (ATV), Decoquinate (DCQ), Tetracyclic Benzothiazepine (TBZ), GW844520 (GW8), GSK932121 (GSK9), Endochin-like quinolones (ELQ), KAE609 (KAE6), GNF-Pf-4492 (GNF4), PA21A092 (PA21), SJ733 (SJ73), AN3661 (AN36), Fosmidomycin (FOSM), MMV008138 (MV81), BMS-388891 (BMS3), MMV019066 (MV19), Pyrimethamine (PYR), Proguanil (PGL), MMV027634 (MV27). Created with Biorender.com

**Figure 8.**
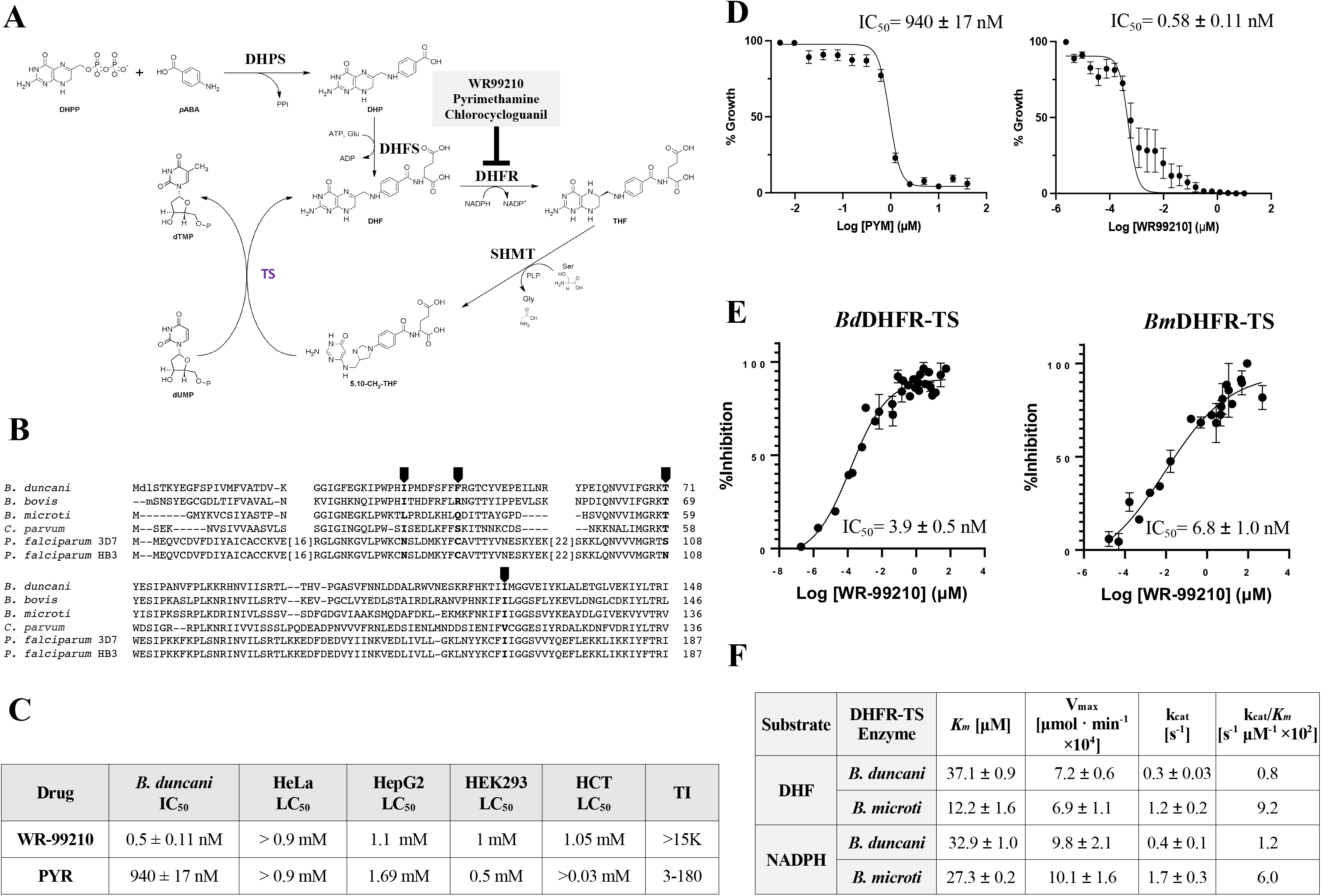
DHFR-TS from *Babesia* parasites as a potential target for therapeutics development. A) Overview of the synthesis of the folate metabolism and inhibitors. Synthesis of dihydrofolate (DHF) precursor by DHP(F)S and reduction of dihydrofolate (DHF) to tetrahydrofolate (THF) catalyzed by the bifunctional enzyme dihydrofolate reductase (DHFR) by using NADPH as electron donor. The thymidylate synthase (TS) catalyzes the reductive methylation of deoxyuridine monophosphate (dUMP) in deoxythymidine monophosphate (dTMP) using the tetrahydrofolate (5,10-Methylene THF) as cofactor via a hydroxymethyl transfer, mediated by serine hydroxy methyltransferase (SHMT). Common DHFR inhibitors are marked in gray box. B) Sequence comparison of DHFR-TS sequences between *B. duncani, B. microti, B. bovis and P. falciparum* pyrimethamine-sensitive (3D7) and-resistant (HB3) parasites with highlighted resistance-associated residues in their primary sequences (black arrows). C) Therapeutic indexes of antifolates against different human cell lines cultured and tested *in vitro.* Each experiment was repeated thrice (*n=3*) for mean ± SD. D) Dose-response curves with IC_50_ values of antifolates pyrimethamine and WR-99210 and their inhibitory effect on *B. duncani* intraerythrocytic parasite growth. E) Enzymatic activity inhibition by WR-99210 and F) kinetic parameters of purified recombinant DHFR-TS enzyme. The rate of a reaction with purified recombinant *Bd* and *Bm* DHFR-TS enzyme against both DHF and NADPH substrates was measured by conducting *in vitro* biochemical assay, and determined the *K_m_*, V_max_ and K_cat_ by a steady-state kinetics using Graph Pad Prism. The IC_50_ values of WR-99210 was determined by plotting a dose response curve using Graph Pad Prism.

## Discussion

In this study, we report the first genome sequence assembly, 3D structure, annotation, transcriptomic profile and metabolic reconstitution of the human intraerythrocytic pathogen *B. duncani*. Recent studies have shown that this parasite is unique among hematozoa, the subclass of apicomplexan pathogens that includes *Babesia*, *Plasmodium* and *Theileria* parasites. The high GC content of this parasite, the ability to propagate continuously in human red blood cells *in vitro* and the availability of a mouse model of virulence make it an ideal system to study the basic requirements for intraerythrocytic parasitism and identify conserved and essential metabolic processes that could be targeted for the development of pan-antiparasitic drugs. Inhibitors of such processes could then be evaluated in mice for safety and efficacy to select the best candidates to advance towards clinical evaluation. Several new drug targets and potential drugs that could inhibit their function have been identified in this study and will stimulate future efforts to evaluate their activity using the *B. duncani* ICIM model system. As a proof of feasibility, we have validated the DHFR-TS enzyme of *B. duncani* and *B. microti* as excellent targets for the development of antibabesial drugs. Using purified recombinant proteins from both pathogens, we showed that pyrimethamine and WR99210 are potent inhibitors of these enzymes. *In vitro* efficacy studied further showed that these drugs block *B. duncani* development in human red blood cells in the nM to sub-nM range. While pyrimethamine is an approved treatment for several parasitic diseases, WR99210, while a more potent drug, has significant hERG liabilities that need to be further addressed before they can be advanced for further preclinical and clinical evaluation.

Our annotation of the *B. duncani* genome provided invaluable insights into the biology, pathogenesis and drug susceptibility of this important human pathogen. Notable among these is the presence of classes of multigene families that could play an important in host-pathogen interactions and virulence of the parasite in mice and humans. The BdOMGF families encompass genes with orthologs in other hematozoan parasites and thus might have evolved to support the survival of the parasite in mammalian red blood cells and escape from host immune attacks as was shown for the var and VESA gene families of *P. falciparum* and *B. bovis* (40).

Our analysis of the *B. duncani* proteome further identified a putative carbonic anhydrase BdGPI09 as a GPI-anchored protein in this parasite, a property also shared with human and mouse CA IV enzymes, which play a critical role in catalysis of reversible CO_2_ hydration and pH regulation of the cytoplasm. Current progress in the development of genetic tools to manipulate *B. duncani* will make it possible to investigate the importance of this enzyme in Babesia survival in human red blood cells and virulence in mice. Noteworthy, while other apicomplexan parasite express CA enzymes, these proteins are not predicted to be GPI-anchored. However, further analyses are needed to determine whether they are membrane associated and whether membrane association is critical for parasite survival in the host cell.

The annotation of the *B. duncani* genome and the analysis of orthology groups have provided insights into the evolution of this parasite among *Apicomplexa*. Binary distance between orthology groups places *B. duncani* at the root of true-*Babesia* when selecting groups with genes in more than three isolates (Fig. S2). A gene orthology-based phylogenetic tree agrees with the proposed evolution of *Babesia* species (Fig. S2). *Babesia microti* was not considered in this comparison as it is distantly related to the main group of piroplasmida. In fact, *B. microti* is part of an early evolving group among the phylum (25). Classification based on OrthoMCL groups shows two clusters encompassing *B. duncani* proteins that were enriched either in Theileridae or Babesidae parasites. The Theileridae specific cluster is composed of 287 IDs among which 73 contain *B. duncani* genes. We identified 29 proteins that were shared with Theileridae and absent from other piroplasmida, including *B. microti* as well as *P. falciparum*. Half of the proteins were of unknown function and nucleotide binding domain was the most predicted function. This sub-group of proteins includes one DNA binding proteins of the AP2 class, one RAD51-like protein, and a putative RNA-binding RAP protein associated with Pfam domain PF08373 that could be targeted to the mitochondria as was shown in *P. falciparum* (41). The Babesidiae cluster is composed of 554 OrthoMCL IDs (Fig. S3), 259 of which are *B. duncani* proteins. 33 proteins were associated with OrthoMCL ID that were unique to true-*Babesia* species. Most functions in this group are related to the regulation of gene or protein expression. BdWA1_000742 was the only surface protein which was part of this group of *Babesia* specific protein. It is a one transmembrane domain protein with four EGF-like domains (hEGF, PF12661) with more divergent homologs in *Plasmodium* and *Theileria*.

Another unique property of *B. duncani* is the presence of a large array of tandemly duplicated functions. We found 139 genes with OrthoMCL ID that had unique genes in other piroplasmida, but have two or up to four genes in *B. duncani*. These genes vary in size and sequence suggesting that these duplications may be ancient and continue to occur in the genome. No specific enrichment of specific functions was observed, but they could altogether contribute to an increase in the metabolic activity of the parasite. Two DNA polymerase subunits and RPB11 subunit of the polymerase II are duplicated as well as TFIIA basal factor and other RNA binding and/or modifying proteins. Duplicated functions were also associated with cellular trafficking, ubiquitinylation and stress response. No enzymes from the central metabolic pathways and subunit of the ribosome were part of this specific group of duplicated genes.

In summary, our analysis of the *B. duncani* nuclear genome and transcriptome has shed light into the biology, evolution, and drug susceptibility of this important human pathogen. It is anticipated that this knowledge will help advance new strategies to develop reliable, sensitive, and specific diagnostic tools as well as therapeutic strategies for a better management of the disease. The close relationship between *B. duncani* and other hemoparasites including malaria parasites will help future efforts to better understand the evolution of virulence in Apicomplexa.

## Material and methods

### Parasite strain

The *B. duncani* WA1 used in this study was obtained from BEI Resources (Cat. No. NR-12311)) and propagated either *in vitro* or in mice and hamsters as previously described (11).

### *In vitro* parasite culture of *B. duncani*

The *in-vitro B.duncani* parasites were cultured in human RBCs as reported earlier (11). Briefly; parasites were cultured in a complete HL-1 medium (Base medium of HL-1 (Lonza, 344017) supplemented with 20% heat-inactivated FBS, 2% 50X HT Media Supplement Hybrid-MaxTM (Sigma, H0137), 1% 200 mM L-Glutamine (Gibco, 25030-081), 1% 100X Penicillin/Streptomycin (Gibco, 15240-062) and 1% 10 mg/mL Gentamicin (Gibco, 15710-072)) in 5% hematocrit A^+^ RBCs the cultures are maintained at 37°C under a 2% O_2_ / 5% CO_2_ / 93% N_2_ atmosphere in a humidified chamber. The culture medium was changed daily and monitored the parasitemia by light microscope examination of Giemsa-stained thin-blood smears.

### Chemicals

Unless otherwise stated, chemicals were purchased from commercial suppliers and used as received. WR99210, JPC2067 and JPC2056 with ≥ 95% pure by reversed-phase high-performance liquid chromatography (HPLC) were purchased form Jacob’s pharmaceuticals company.

### DNA prep for PacBio sequencing

Genomic DNA was isolated from 100 ml *in vitro* culture of *B. duncani* (15% parasitemia and 5% hematocrit) using DNasy Blood and Tissue kit (Qiagen; Cat. No. 69506), and quality control along with concentration determination was performed using nanodrop and qubit. DNA integrity was evaluated using Blue Pippin pulse gel and the DNA was then used for library preparation using Pacific Biosciences SMRTbell Express template Prep Kit 2.0 (Cat. No. PN: 100-938-900) according to the manufacturer’s instructions. The Pacific Biosciences Smart Link software was used to determine loading concentration and proper stoichiometric measurements. The gDNA library was then annealed to the Pacific Biosciences V5 primer for 1h at 20°C. The annealed library was then bound to polymerase using Pacific Biosciences Polymerase 2.2 for 1-4h at 30°C and was loaded on to the Sequel II Instrument as an adaptive sequencing run. At least one smart cell was sequenced for each genomic DNA library with a movie time of 30h and a pre-extension of 2h. After the DNA library sequencing was complete, the loading metrics were evaluated by mean read length, polymerase read length, data yield and P1 values to ensure the sample ran as expected and data had met Yale’s gold standards.

### DNA preparation for Bionano optical map

Exactly 3 ml packed frozen pellets of *B. duncani* WA1 in human RBCs were used to isolate ultra-high molecular weight (uHMW) genomic DNA for use in genomic optical mapping by Histogenetics (Ossining, NY) using the Bionano Prep™ Blood and Cell Culture DNA Isolation Kit (Bionano Genomics, cat No. 80004). Following this, DNA was quantified using Qubit™ dsDNA BR Assay Kit. A total of 0.75 ug of HMW DNA was then labeled using the Bionano Prep direct label and stain (DLS) method (Bionano Genomics, cat No. 80005) and loaded onto a flow cell to run on the Saphyr optical mapping system (Bionano Genomics). Approximately 1,177 Gb of data was generated per run. Raw optical mapping molecules in the form of BNX files were run through a preliminary bioinformatic pipeline that filtered out molecules less than 150 kb in size with and less than 9 motifs per molecule to generate a *de novo* assembly of the genome maps.

### Genome sequencing and assembly

100 mL *in vitro* culture of *B. duncani* in hRBCs was propagated to a parasitemia of 10% at 5% HC. DNA was isolated from infected hRBCs and sequenced at the Yale Center for Genome Analysis using Illumina HiSeq 2500 and PacBio HiFi (CCS) sequencing. A total of 177,189 PacBio HiFi reads were obtained; the average HiFi read length was 8,814 bp; the longest read was 29,067 bp. The HiFi reads totaled 1.56B bases which translated to an expected ∼156x coverage of the *B. duncani* genome (assuming a ∼13Mb genome). HiFi reads were mapped using Minimap2 (42) to the *B. duncani* mitochondrion and apicoplast. Only 0.41% of the reads mapped to these organelles, which were then discarded to enrich the data for nuclear DNA. HiFi reads were also mapped to the human genome to determine possible host contaminations, but only 1.48% of them were flagged. Cleaned reads were assembled using HiCANU v2.2 (18) using default parameters. HiFiASM v0.16 (43) and Wengan (44) were also tested on the HiFi reads, but they generated slightly inferior assemblies (based on assembly statistics, comparison with the optical map and transcript isoform annotations). The Bionano Hybrid Scaffolding pipeline v1.7 was used to compare the draft assembly with the Bionano optical map to detect possible mis-joins and to create scaffolds. The scaffolded assembly was polished using two rounds of PolyPolish (19). At every step of the assembly (draft, scaffolds, polished), a series of quality control steps were carried out to ensure that no imperfections were introduced. The QC steps included (1) mapping all the Illumina WGS and HiFi reads to the assembly, (2) carrying out a BUSCO (23) completeness analysis (at the genome level), (3) determining the number of detected gene loci from the mapping of RNA-Seq IsoSeq reads.

### DNA preparation for Hi-C

100 ml *in vitro* culture of *B. duncani* WA1 in A^+^ human RBCs was grown to a parasitemia of 15%. The culture was centrifuged, and the parasite pellet was crosslinked in 1.25% formaldehyde for 25 min at 37°C. Crosslinking was quenched in a final concentration of 150 mM glycine for 15 minutes at 37°C followed by a 15-minute incubation at 4°C. Parasite pellets were then resuspended in lysis buffer (10 mM Tris-HCl, pH 8.0, 10 mM NaCl, 2 mM 4-(2-aminoethyl) benzenesulfonyl fluoride HCl (AEBSF), 0.25% Igepal CA-360 (v/v), and EDTA-free protease inhibitor cocktail (Roche)) and incubated for 30 min on ice. Nuclei were isolated after homogenization by 15 needle passages. *In situ* Hi-C protocol was performed as described by Rao and colleagues (45). Briefly, nuclei were permeabilized using 0.5% SDS. DNA was digested with 100 units of Mbol (NEB), the ends of restriction fragments were filled using biotinylated nucleotides and ligated using T4 DNA ligase (NEB). After reversal of crosslinks, ligated DNA was purified and sheared to a length of ∼300-500 bp using the Covaris ultrasonicator S220 (settings: 10% duty factor, 200 cycles per burst and a peak incident power of 140). Ligated fragments were pulled down using streptavidin beads (Invitrogen) and prepped for Illumina sequencing by subsequent end-repair, addition of A-overhangs and adapter ligation. Libraries were amplified for a total of 12 PCR cycles (45 sec at 98°C, 12 cycles of 15 sec at 98°C, 30 sec at 55°C, 30 sec at 62°C and a final extension of 5 min at 62°C) and sequenced with the NOVASeq platform (Illumina), generating 100 bp paired-end sequence reads at the UCSD core facility.

### Hi-C data processing

The Illumina sequencing of the four *B. duncani* clones (WA1, A6, B7 and B1) yielded over one billion Hi-C paired-end reads in total. Hi-C reads were processed using the command-line version of the HiCExplorer pipeline (46). HiCExplorer is a comprehensive and versatile pipeline which processes Hi-C reads and generates normalized chromatin conformation contact maps. The pipeline started by mapping all Hi-C single reads to the *B. duncani* assembly using “BWA mem -A1 -B4 -E50 -L0” (47). The percentage of reads mapped ranged between 83% and 96%. After sorting and merging the reads from the four data sets, the command “hicBuildMatrix” was used with default parameters, and a bin size of 10Kb. A diagnostic plot created using “hicCorrectMatrix diagnostic_plot” indicated that a threshold of -4.5 would be appropriate to remove GC and open chromatin biases. The correction step, carried out by the command “hicCorrectMatrix correct”, also normalized the number of restriction sites per bin. The contact maps were plotted using “hicPlotMatrix”, as part of the same pipeline.

### 3D modeling

The 3D model of the *B. duncani* genome was generated and visualized using PASTIS (48) and ChimeraX (49). Briefly, interaction data were first manually fitted to a three-column interaction count matrix, where the first two columns indicate the two separate bins within the genome and the third being the number of interactions between those two bins. Then the three-dimensional coordinate matrices were generated using PASTIS and converted to .PDB format, with each line being one of the coordinates output by Pastis. The data was visualized using ChimeraX (49).

### PacBio IsoSeq processing

PacBio IsoSeq reads were mapped to the *B. duncani* genome using Minimap2 (42) with options “splice:hq -uf --secondary=no -C5”. The resulting SAM files sorted using “sort -k 3,3 -k 4,4n” then fed into the PacBio cDNA_Cupcake ToFU pipeline (https://github.com/Magdoll/cDNA_Cupcake) using the Python code “collapse_isoforms_by_sam.py” to collapse redundant isoforms (more details about this pipeline can be found at https://github.com/Magdoll/cDNA_Cupcake/wiki). The resulting isoform sequences were used in the gene finding pipeline below.

### Gene prediction and annotation

The genome was first soft-masked with RepeatMasker (http://www.repeatmasker.org), then processed using the FunAnnotate v1.8.9 gene annotation pipeline (https://github.com/nextgenusfs/funannotate). As transcript evidence, we provided the IsoSeq-based isoforms computed via the cDNA_cupcake pipeline (see PacBio IsoSeq processing). As protein sources, we provided the annotated protein sets of *B. bigemina*, *B. bovis*, *B. microti*, *P. falciparum*, *T. gondii*, *T. orientalis*, *T. parva* as well as all the UniProt/SwissProt protein models. FunAnnotate was ran with default parameters and weights (augustus:2 hiq:4 transcripts:4 proteins:4). Functional annotations were obtained via InterProScan v5.55-88 with default parameters (50). The output of InterProScan was parsed by custom scripts for the downstream analyses.

### Synteny and gene localization plots

Synteny plots were obtained using mummer2circos (https://github.com/metagenlab/mummer2circos) with the promer algorithm, and Circos (51). Gene localization plots were produced using custom Python scripts and the Biopython Bio.graphics library.

### Orthology detection and database searches

*B. duncani* genes were assigned to OrthoMCL (https://OrthoMCL.org) groups using the orthology assignment tool available through the VEuPathDB (https://VEuPathDB.org) Galaxy workspace (52, 53). 4,170 Translated *B. duncani* proteins in FASTA format were assigned to groups based on the OG6r9 blast database using the default settings. Output files generated by the OrthoMCL pipeline included a mapping file between *B. duncani* gene IDs and OrthoMCL v.6 group IDs (this file was used to query the OrthoMCL database to determine degrees of evolutionary conservation) and a file of leftover proteins that did not map to any OrthoMCL groups but did cluster with at least one other *B. duncani* protein (these were considered species specific gene duplications or families). To identify genes that do not have any orthologs or paralogs, the original input FASTA file was parsed for *B. duncani* IDs that are not present in any of the OrthoMCL mapping output files. VEuPathDB resources including PlasmoDB.org, ToxoDB.org, CryptoDB.org and PiroplasmaDB.org were used to retrieve genome size, gene content and chromosome numbers.

A matrix containing the number of genes per OrthoMCL groups was made from the annotation of *B. duncani* and species present in PiroplasmaDB. Annotations were compared by selecting OrthoMCL groups presenting annotation in at least 4 species. *B. microti* reduces drastically the number of OrthoMCL IDs and was removed from the analysis. OrthoMCL IDs were compared using Euclidian distance and hierarchical classification was performed using Ward method. Species annotation were compared using presence/absence of the OrthoMCL ID. Jaccard distance and Ward methods were used to make the tree. All analyses were performed in R. The ComplexHeatmap function was used to generate the heatmap.

### RNA-Seq processing for gene-expression analysis

RNA-seq FASTQ files were assessed for quality using FastQC (version 0.11.8). Adapter sequences as well as the first 11 bp of each read were trimmed using Trimmomatic (version 0.39). Tails of reads were trimmed using Sickle with a Phred base quality threshold of 25, and reads shorter than 18 bp were removed. Reads were then aligned to the *B. duncani* genome assembly using HISAT2 (version 2.2.1). Only properly paired reads with a mapping quality score of 40 or higher were retained, with filtering done using Samtools (version 1.11). StringTie (version 2.2.1) was run with the -e parameter to estimate the abundance of each gene in TPM (transcripts per million).

### Three-dimensional (3D) modeling and visualization of *B. duncani* genome architecture

To investigate the spatial conformation of the *B. duncani* genome, a 3D model of the three chromosomes was first built using PASTIS (48) and then improved using the Bezier curve smoothing method. PASTIS infers a consensus 3D structure from the genome-wide Hi-C contact frequency matrix using the following probability model. It models the observed chromatin contact frequencies as independent Poisson random variables and infers the model parameters as well as the 3D coordinates of the genome structure via the maximum likelihood estimation approach. Specifically, PASTIS assumes the Poisson parameter λ_ij_for chromatin interaction frequency c_ij_ between loci i and j as a decreasing function of d_ij_ (11),

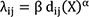

with parameters β > 0 and α < 0. Here d_ij_ (11) (or simply d_ij_) is the Euclidean distance between loci i and j in structure X. Therefore, the likelihood of observing c_ij_’s can be formulated as:

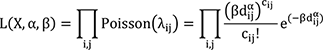

By maximizing the log-likelihood function, PASTIS optimizes the 3D structure X as well as the distance decay parameters α and β.

Since the 3D model predicted by PASTIS contained a set of discrete points, cubic Bezier curve smoothing was applied for visualization purpose. Cubic Bezier curves guarantees the interpolated curve passes through all points in the original structure and the first-order geometric continuity at all points are maintained.

### GPI-associated proteins (GPI-AP) identification

GAPI-AP were selected based on a two transmembrane domain model, SignalP 5.0 and PredGPI online predictions. TMPred software was used to validate the presence of hydrophobic regions associated with the N-terminal signal peptide (ER-targeting) and C-terminal GPI-anchor signal (recognized by the transamidase). The maximum distance between the N-terminal TM with the start of the sequence was set to 10 with a minimal score of 500. The maximum number of residues between the C-terminal TM with the end of the sequence was 3 with a minimal score of 780. Internal hydrophobic helix should not have score higher than 800. We keep protein that were positive for the three predictors. Proteins lower than 100 aa were not considered.

### Drug efficacy study with *in vitro B. duncani*-infected human red blood cells

The effect of antifolates on intra-erythrocytic development cycle (IDC) inhibition of *B. duncani* and the IC_50_ determination was done by the reported protocol (*14*). Briefly, *in vitro* parasite culture (0.1% parasitemia with 5% hematocrit in complete HL-1 medium) was treated with decreasing concentrations of the compound of interest in a 96-well plate for 60 Hrs. After the treatment, parasitemia was enumerated by SYBR Green-I method by adding equal volumes of parasite culture with lysis buffer (0.008% saponin, 0.08% Triton-X-100, 20 mM Tris-HCl (pH = 7.5) and 5 mM EDTA) containing SYBR Green-I (0.01%) and incubated at 37°C for 1h in the dark. The fluorescence was measured at 480 nm (excitation) and 540 nm (emission) by using a BioTek Synergy™ Mx Microplate Reader. Fluorescence from the background (uninfected RBCs in complete HL-1medium) was subtracted from each concentration and determined the 50% inhibitory concentration (IC_50_) of the drug by plotting sigmoidal dose-response curve fitting with drug concentration and per cent parasite growth in the Graph Pad prism 9.2.1 from two independent experiments with biological triplicates. Data are shown as mean ± SD.

### Assessment of drug cytotoxicity on human cell lines

We obtained HeLa, HepG2, HEK and HCT116 cell lines from the American Type Culture Collection and maintained them in Dulbecco’s modified Eagle’s medium (DMEM) (Invitrogen 11995-065) containing 25 mM glucose, 1 mM sodium pyruvate and supplemented with 5 mM HEPES, 10% FBS and penicillin-streptomycin (50 U/mL penicillin, 50 µg/mL streptomycin). Seeded 20,000 cells per well in a 96 well tissue culture plate and allowed to adhere. After 24 hours, cells were treated with 2-fold serially diluted drugs [from 10 mM as highest final concentration] and parallelly with the 0.1% and 10% DMSO [as negative and positive vehicle controls respectively] and incubated at 37°C for 72 hours. After the drug treatment, each well was incubated with 0.5 mg/ml of MTT reagent [Cayman chemical Cat# 10009591] for 4 hours in the dark at 37°C. The formed formazan crystals by living cells were solubilized with the addition of 100µl of DMSO to each well and measured the OD of samples at 590 nm using Spectra max plate reader. From the obtained OD values, per cent cell viability by normalizing to the mean of 10% DMSO wells (set as 100% toxicity) and mean of the vehicle control wells (set as 0% toxicity). Dose-response curves were plotted using GraphPad Prism version 9.1.2

### Steady-state kinetics of DHFR-TS activity

DHFR kinetic experiments were performed by incubating purified enzyme (0.1 μM) with a saturating concentration of DHFR ligand (NADPH or DHF at 300 μM) and measuring the reaction rate at varying concentrations of the complementary ligand (DHF or NADPH, respectively). Experiments were performed in a 96-well plate with reaction buffer: 25 mM Tris pH 8, 20 mM NaCl, 50mM L-arginine, 0.5% glycerol, 1 mM DTT and 1 mM ethylenediaminetetraacetic acid (EDTA). Recombinant DHFR activity was measured by a decrease in absorbance at 340 nm as NADPH was converted to NADP+. Plates were incubated at 37°C, and optical density (OD_340nm_) measurements were taken using a BioTek SynergyMx microplate reader for 1 min. Rate constants for steady-state kinetic experiments were estimated by fitting the data to a Michaelis−Menten hyperbolic curve (v = Vmax[S]/(Km + [S])), where v is the reaction rate, [S] the concentration of substrate, and Km the Michaelis constant) using the curve-fitting program, using GraphPad Prism.

### The half-maximal inhibitory concentration (IC_50_) of DHFR enzymatic activity

The inhibition of selected compounds on DHFR activity was measured by incubating purified enzyme (0.1 μM) with rising concentration of drugs. The reaction buffer contained 300 μM DHF and 300 μM NADPH, and the OD_340nm_ reduction rate (OD_340nm_/min) was documented for further calculation. The % inhibition was calculated and normalized to DMSO (100% activity) and no enzyme (0% activity) wells accordingly: 1-[((OD_340nm_/min)_sample_-(OD_340nm_/min)_neg. control_))/(OD_340nm_/min)_pos. control_-(OD_340nm_/min)_neg. control_))]. The IC_50_ was determined from a sigmoidal dose-response curve using GraphPad Prism version 9.1.2.

## Acknowledgements

We thank Ruiyi Gao for her contribution to the initial efforts to sequence the *B. duncani* genome. C.B.M.’s research is supported by grants from the National Institutes of Health (AI097218, GM110506, AI123321 and R43AI136118), the Steven and Alexandra Cohen Foundation [Lyme 62 2020], and the Global Lyme Alliance. and the Global Lyme Alliance. SL’s research is supported by grants by the US National Science Foundation (IIS 1814359) and the National Institutes of Health (1R01AI169543-01). KLR’s research is supported by the National Institutes of Allergy and Infectious Diseases (R01 AI136511, R01 AI142743-01 and R21 AI142506-01), the University of California, Riverside (NIFA-Hatch-225935) and the Health Institute Carlos III (PI20CIII/00037).

## Conflict of interest

- There are no actual or perceived conflicts of interest on the part of any author
- The authors declare that they have no conflicts of interest with the contents of this article

---

**Figure S1:**
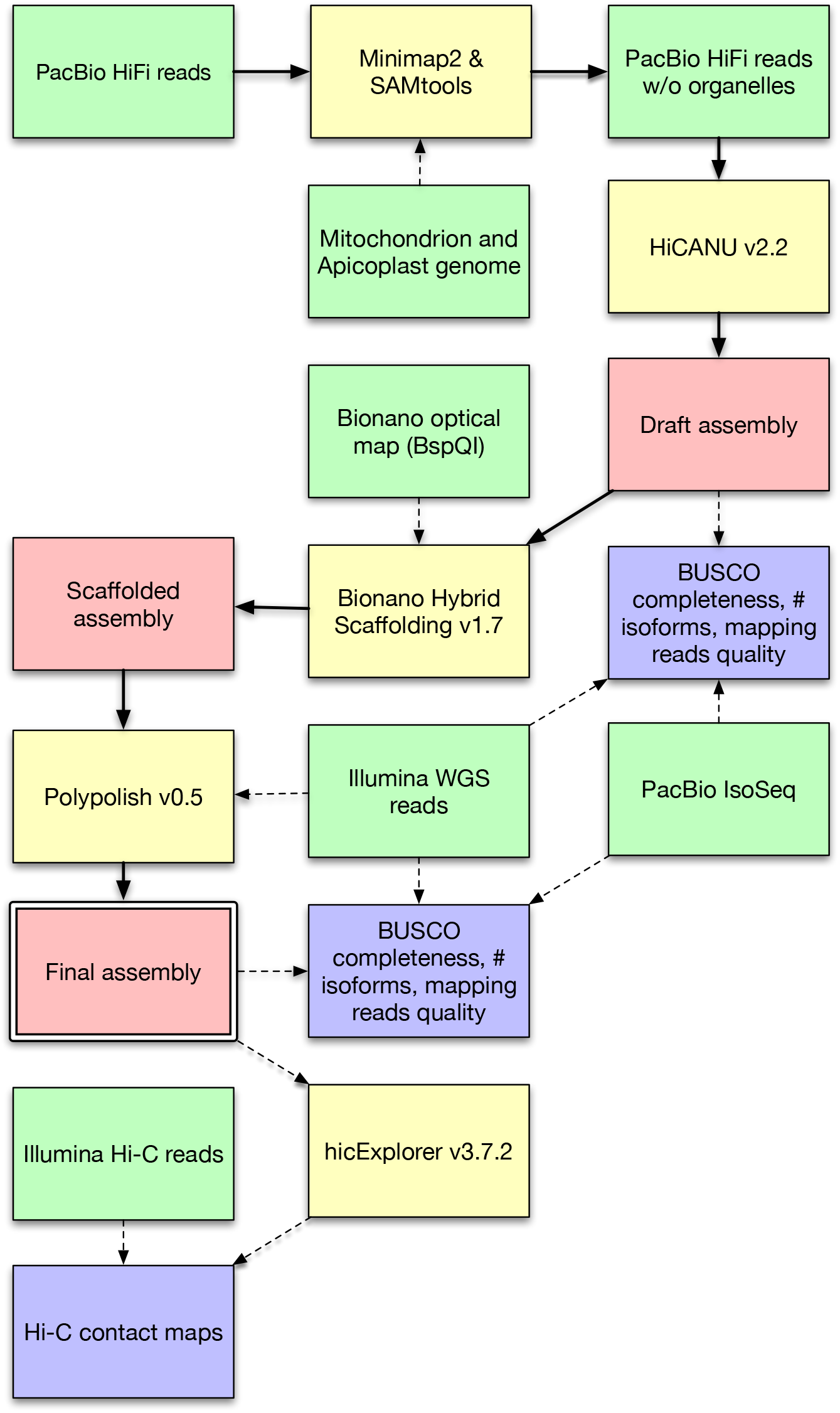
Software pipeline for the assembly of the *B. duncani* genome. Green blocks indicate genomics data sets; blue blocks indicate quality control terminal points; yellow blocks indicate software tools

**Figure S2:**
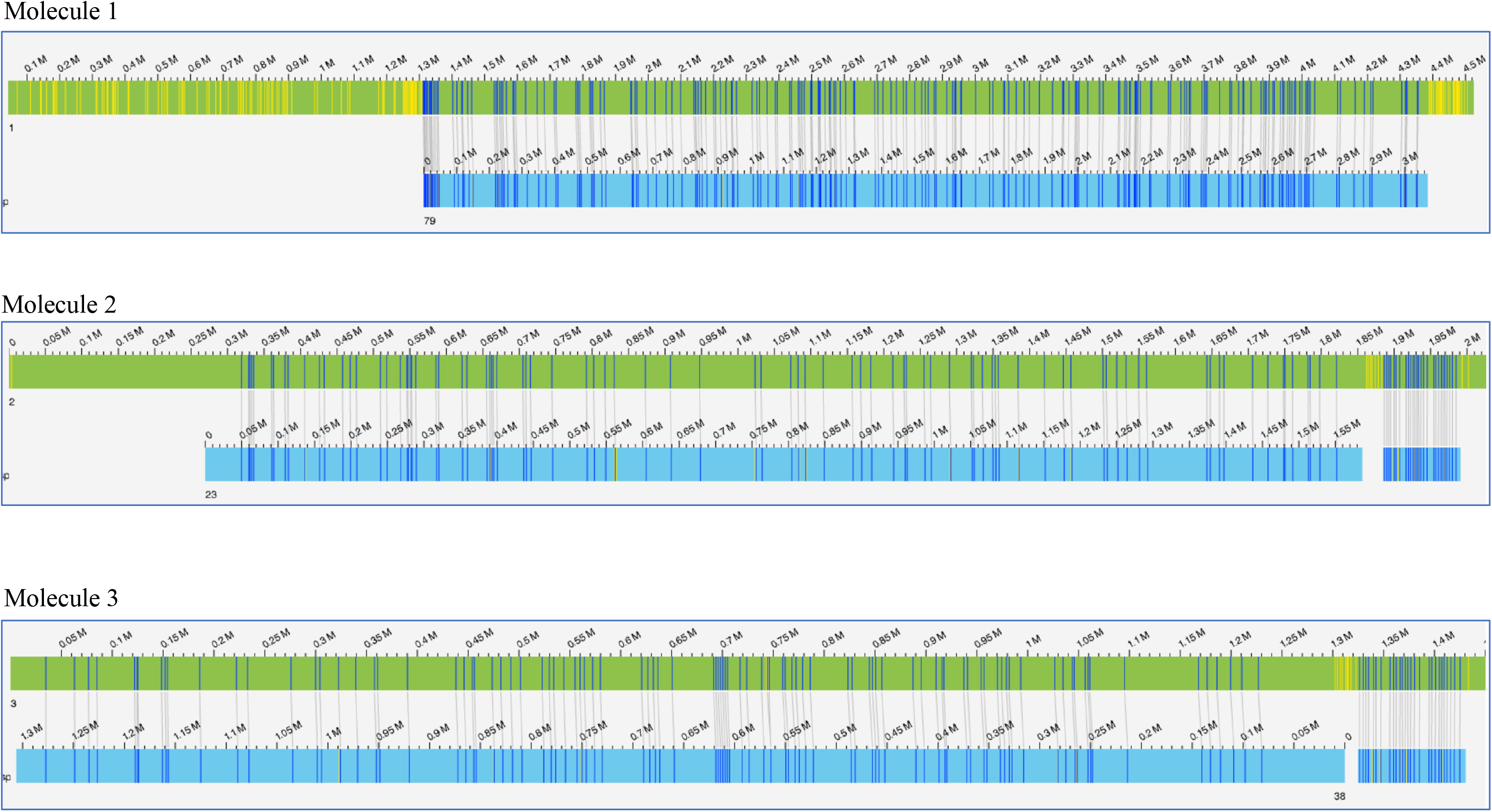
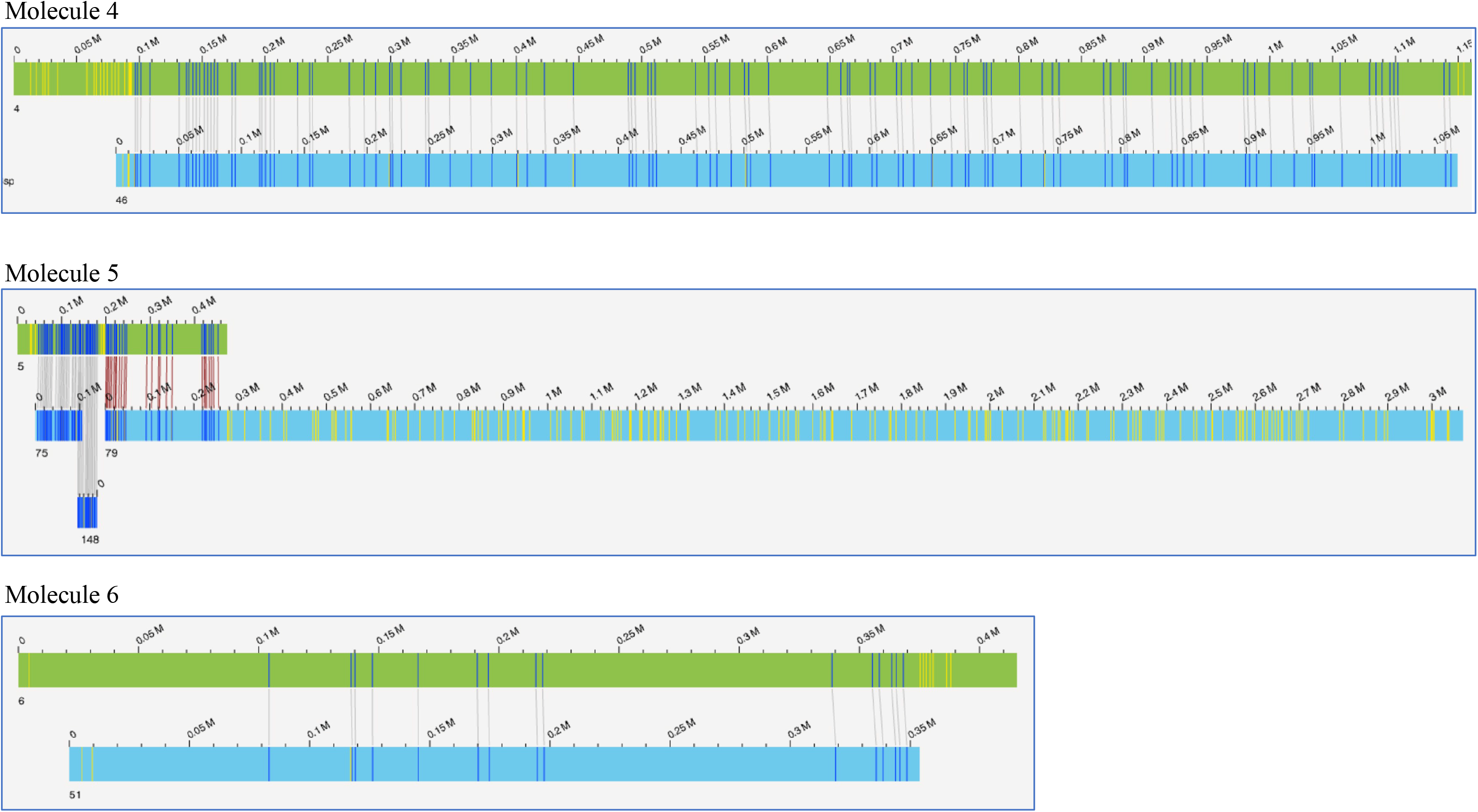
*B. duncani* OrthoMCL annotation and Babesia evolution. **A.** OrthoMCL annotation is summarized in a heatmap format. Groups were clustered based on the number of present genes. Grey color was used when no genes were present in a species. Light red means that only one gene is present and dark red correspond to OrthoMCL groups presenting more than 1 gene. Two clusters were specific of Theileridae on one side and true-*Babesia* on the other side. The top tree is the same as in B. **B.** Evolution of piroplasmida based on OrthoMCL annotation. The Jaccard distance was used for binary analysis of the presence/absence of OrthoMCL groups in each species.

**Figure S3.**
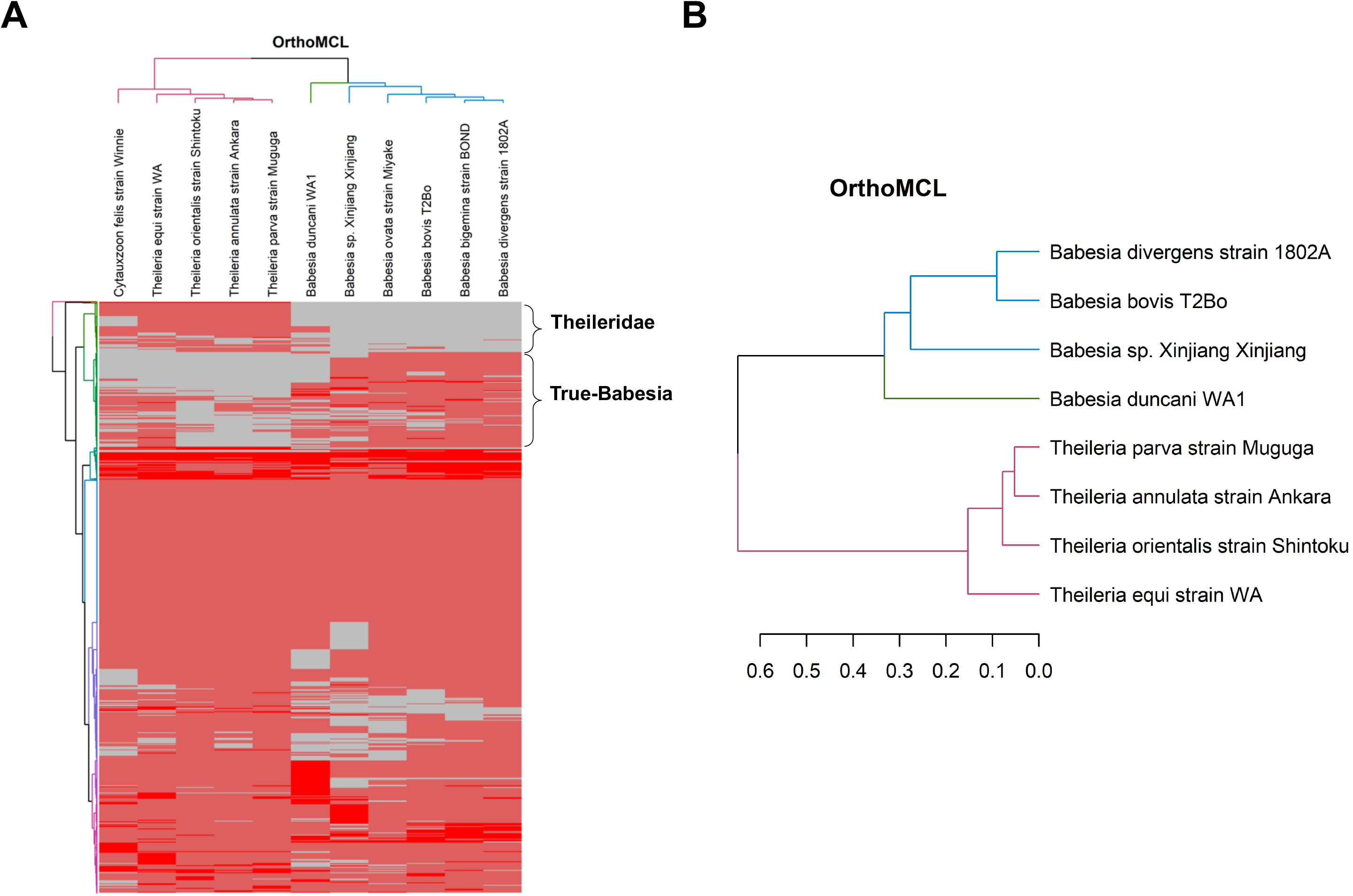
Optical Map (1/2). Alignment of the *Babesia duncani* genome assembly against molecule 1, 2 and 3 of the *B. duncani* optical map (BspQI).

**Figure S4.**
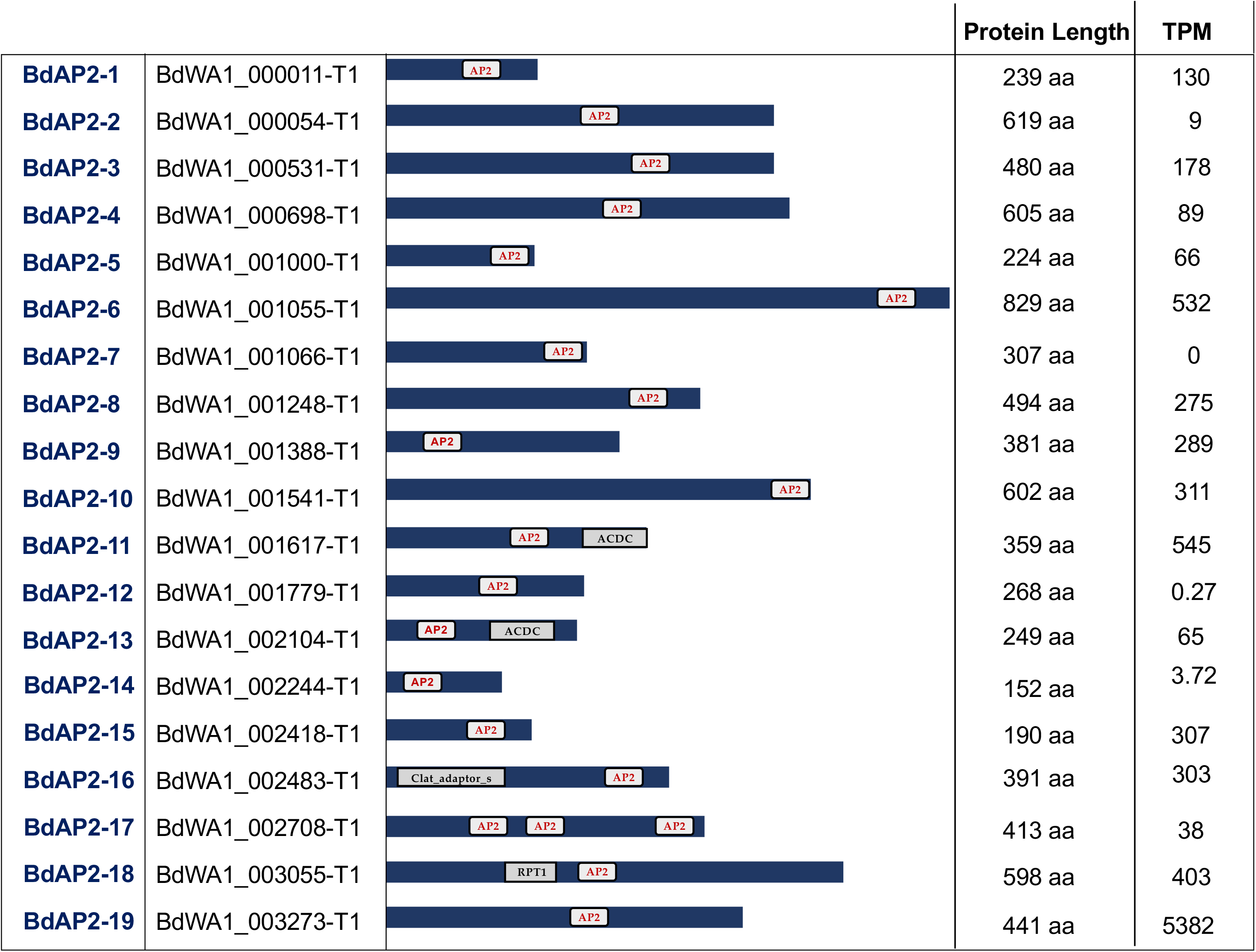
Optical Map (2/2). Alignment of the *Babesia duncani* genome assembly against molecule 4, 5 and 6 of the *B. duncani* optical map (BspQI).

**Table S1:** Length of the six molecules in the *Babesia duncani* optical map (BspQI).

| BspQI | Length (bp) |
| --- | --- |
| 1 | 4,525,065 |
| 2 | 2,026,521 |
| 3 | 1,450,238 |
| 4 | 1,160,926 |
| 5 | 474,067 |
| 6 | 415,637 |
| SUM | 10,052,454 |

**Table S2.**
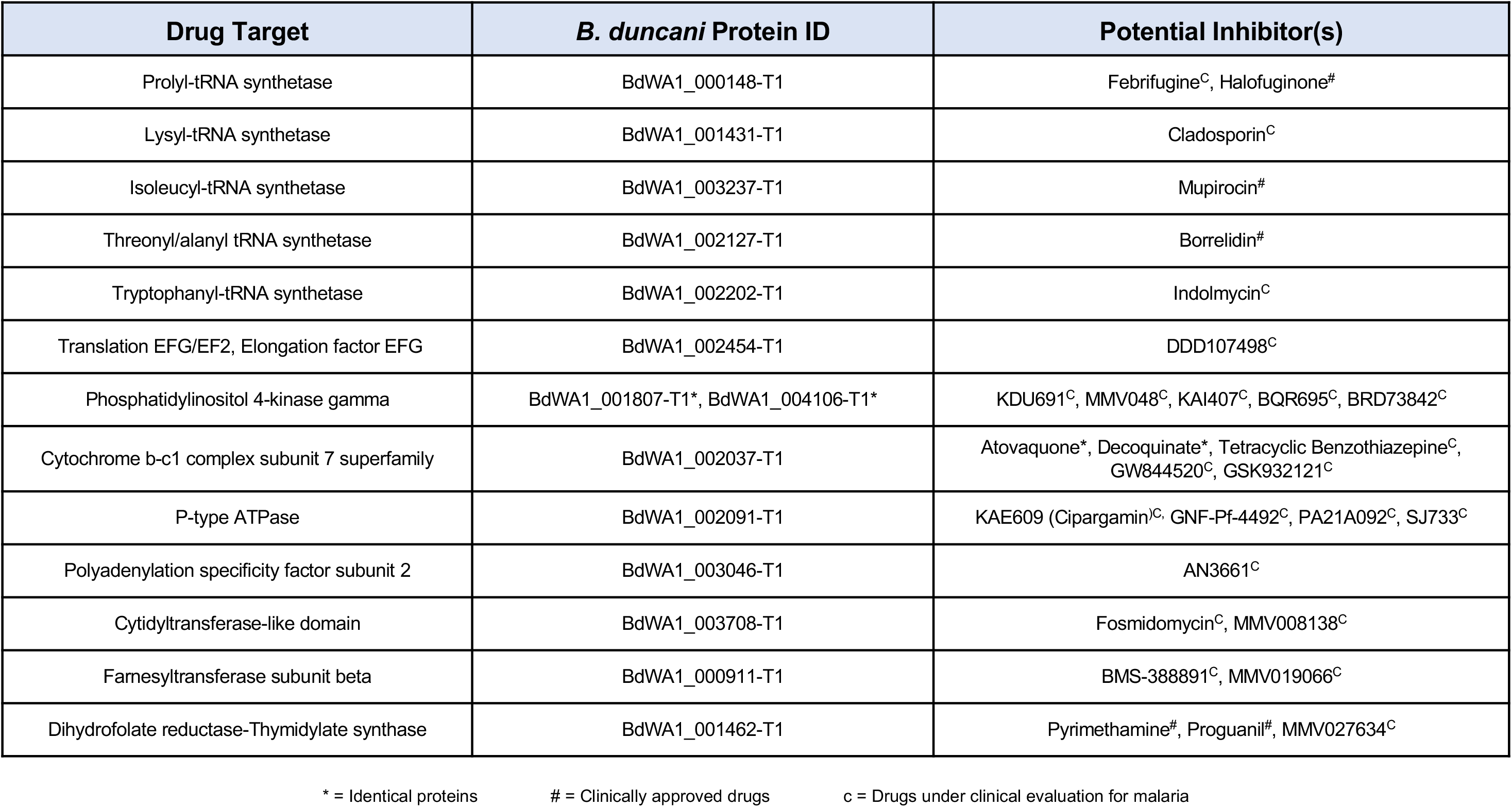
Emerging drug targets for human Babesiosis therapy.

## Supplemental Methods

### RNA preparation for Illumina RNA-Seq

Total RNA was isolated from parasites using 5 volumes of 37^°^C pre-warmed Trizol LS Reagent (Life Technologies, Carlsbad, CA, USA) and incubated at 37^°^C for 5 minutes. Total RNA isolation was then continued according to the manufacturer’s instructions. Total RNA was then treated with DNA-free DNA removal kit (ThermoFisher; AM1906) followed by mRNA purification using NEBNext Poly(A) mRNA Magnetic Isolation Module (NEB, E7490S). RNA-seq library was constructed using NEBNext Ultra II RNA-library preparation kit (NEB, E7770S) according to the manufacturer’s instructions. The libraries were amplified for 15 PCR cycles (45s at 98°C followed by 15 cycles of [15s at 98°C, 30s at 55°C, 30s at 62°C], 5 min 62°C). Libraries were sequenced at 150 bp paired-end sequenced on the Illumina Novaseq platform (Illumina, San Diego, CA) at the UCSD and Yale core facility.

### RNA preparation for PacBio Iso-Seq

TRIzol reagent (Life Technologies, Carlsbad, CA, USA, No. 15596–026) was used to isolate total RNA from 100 ml *in vitro* culture of *B. duncani* (15% parasitemia and 5% hematocrit) according to the manufacturer’s protocol. 1 µg of total RNA was used for the synthesis and amplification of cDNA using a combination of NEBNext Single Cell/Low Input cDNA Synthesis & Amplification module (Cat. No. E6421S), NEBNext High-Fidelity 2X PCR Master Mix (Cat. No. M0541S), Iso-Seq Express Oligo Kit (Cat. No. PN 101-737-500), and elution buffer (Cat. No. PN 101-633-500). SMRTbell libraries were constructed according to the Iso-Seq Express Template Protocol (Pacific Biosciences). Primer annealing and polymerase binding were performed following the SMRT Link v8.0 Sample Setup instructions and 90 pM of the SMRTbell templates were loaded for sequencing. One SMRT Cell 8M was used for each sample and sequencing was performed using the Sequel II system.

### Phylogenetic tree reconstruction

To determine the phylogenetic placement of *B. duncani* among a selected group of apicomplexa, a conserved helicase from OrthoMCL group OG6_100305 (PMID: 26738725) was used for the alignments. Proteins from the group that were most similar to *B. duncani* gene BdWA1_002520 were selected and aligned using the MUSCLE (1). Evolutionary history was inferred using the Neighbor-Joining method (2). Bootstrapping analysis was conducted 1,000 times to determine phylogenetic tree reliability. Evolutionary analysis was conducted using MEGA X software (3).

### Pulse field gel electrophoresis (PFGE)

*B. duncani* WA1 cultures were centrifuged at 1300 x g for 5 min to yield pellets containing intact cells. Pellets, were embedded in 1% (w/v) SeaKem Gold Agarose (Lonza, Rockland, ME, USA) to an approximately concentration of 1×10^8^ infected RBCs/ml. The resultant agarose plugs were incubated in lysis solution (100mM EDTA, pH8.0, 0.2% sodium deoxycholate, 1% sodium lauryl sarcosine) supplemented with 1 mg/ml of proteinase K (Thermo Fisher Scientific, Vilnius, Lithuania) for 24 h at 50°C. Finally, plugs were washed 4 times for 30 min each in wash buffer (20 mM Tris, pH 8.0, 50 mM EDTA). Intact chromosomes were separated on a 0.8% Megabase Agarose gel (Bio-Rad Labs Inc., Hercules, CA, USA) in 1X TAE buffer chilled at 14°C for 48 h on a CHEF Mapper^TM^ XA pulsed field electrophoresis system (Bio-Rad). The switch time was 500 sec at 3V/cm with an include angle of 106⁰. The agarose gel was stained with GelRed (Biotium, Fremont, CA, USA) and visualized under ultraviolet.

### Southern Blot Analysis

Telomeric ends of *B. duncani* WA1 chromosomes were analyzed by Southern blot using a nucleotide repeat sequence (CCCTGAACCCTAAA) of the telomeric ends of *Plasmodium berghei* chromosomes (4, 5). The telomeric probe was labeled using the DIG Oligonucleotide Tailing Kit, 2^nd^ Generation (Cat. No. 03353383910, Roche, Mannheim, Germany). After PFGE and before transfer, DNA from the gel was depurinated (20 min in 0.25 M HCL), denatured (2 × 20 min in 0.5N NaOH; 1.5 M NaCl) and neutralized (2 × 20 min in 0.5 M Tris-HCl, pH 7.5; 1.5 M NaCl). Southern blotting was done on nylon membrane, positively charged (Cat. No. 1417240, Roche) using 10X SSC and followed by UV crosslinking of transferred DNA. The membrane was hybridized overnight at 26⁰C with the telomere probe and then washed twice for 5 min each in 2X SSC and for 20 min each in 0.5X SSC at 26°C. Bound probe was detected with disodium-2-chloro-5(4 methoxyspiro (1,2-dioxetane-3.2’-[5-chloro]tricycle[3.3.1.1.^3.7^] decan)-4-yl)-1-phenyl phosphate (CDP-StarTM, Cat. No.12041677001, Roche) according to the manufacturer’s instructions. The membrane was visualized using an Amersham ImageQuant 800 system (GE Healthcare Bio-Science AB, Uppsala, Sweden).

### Cloning, Expression, and Purification of recombinant *B. duncani* and *B. microti* DHFR-TS enzymes

Dihydrofolate reductase-thymidylate synthase (DHFR-TS) gene from *B. duncani* (*Bd*DHFR-TS) and *B. microti* (*Bm*DHFR-TS) were cloned into pMAL-c4x by GenScript USA Inc. Both enzymes were expressed as fusion proteins with N’-terminal maltose-binding protein (MBP) tag. Constructs of pMAL-c4x-*Bd*DHFR-TS and pMAL-c4x-*Bm*DHFR-TS were transformed by heat shock into *E. coli* BL-21(DE3). *E. coli* BL-21(DE3) harboring pMAL-c4x-DHFR-TS was grown overnight in 2 ml LB ampicillin (100 µg/mL). Cultures were diluted 100-fold in fresh LB medium with 0.2% glucose and grown to OD_600nm_ of ∼0.5 at 37°C. DHFR-TS expression was induced by the addition of 0.5 mM isopropyl thiogalactoside (IPTG) followed by growth for overnight at 16°C or at room temperature (RT). The cells were harvested by centrifugation (8,000 × g × 10 min, 4°C), washed by resuspension in water, and re-centrifuged. Cells were used directly for purification or kept frozen at -20°C. Prior to enzyme purification, cells were re-suspended in binding buffer containing 25 mM Tris-HCl pH 8, 500 mM NaCl, 0.5% glycerol, and 50 mM L-arginine. Cells were supplemented benzonase DNAse (250U/µl) and 3-((3-cholamidopropyl) dimethylammonium)-1-propane sulfonate (CHAPS) (0.002%) and were disrupted by sonication on ice using Omni Sonic Ruptor 400 Ultrasonic Homogenizer (15-sec burst at 70% amplitude, 3 times, with 30-sec cooling intervals). A soluble supernatant was prepared by centrifugation (16,000 × g × 20 min) of the cells sonicate.

Recombinant DHFR-TSs were purified from the clear cell extract supernatant using amylose resin (NEB), methotrexate-agarose beads, and in some cases, size exclusion chromatography (SEC). Briefly, the soluble supernatant was incubated with amylose beads in binding buffer with gentle agitation for 1h at 4°C. Following enzyme adsorption, the affinity matrix was transferred to a column, and unbound proteins were removed by washing with a binding buffer. Purified enzymes were eluted with elution buffer containing 10 mM maltose. Amylose elution fractions were loaded onto MTX-agarose resin, and purified DHFR-TS enzymes were eluted using 2 mM dihydrofolic acid (DHF). Maltose or DHF residuals were removed by dialysis or using a PD-10 column (GE Healthcare), and the Purified protein stocks were then adjusted to 25% glycerol v/v, aliquoted, flash-frozen, and stored at -80°C.

## Notes

### Competing Interest Statement

The authors have declared no competing interest.

